# Phage CRISPR-like regulatory RNAs silence multilayered bacterial immune systems

**DOI:** 10.1101/2024.10.29.618823

**Authors:** Chao Liu, Zhihua Li, Rui Wang, Wenhe Wang, Jiachen Yao, Junyuan Xue, Xian Shu, Jing Xu, Guangyi Liang, Feiyue Cheng, Xiaoyu Du, Linlin Guan, Mario Rodríguez Mestre, Jun Du, Lei Cai, Eugene V. Koonin, Yue Feng, Rafael Pinilla-Redondo, Ming Li

**Author notes:** These authors contributed equally to this work (Chao Liu, Zhihua Li, Rui Wang, Wenhe Wang). Corresponding authors. (M.L.); (R.P.-R.); (Y.F.).

## Abstract

CRISPR RNAs (crRNAs) encoded by CRISPR arrays direct Cas nucleases against viruses. Some CRISPR-Cas loci also contain mini-arrays encoding crRNA-like RNAs (crlRNAs), which modulate transcription of *cas* or associated toxin genes that safeguard CRISPR-Cas activity. Here, we report that phages hijack these host crlRNA regulatory circuits to suppress bacterial immunity. Viral crlRNAs compete with host crRNAs for Cas protein binding and reprogram Cas proteins to silence their own expression or suppress transcription of CRISPR-safeguarding toxins. These anti-defense crlRNAs are found across diverse phages and plasmids, frequently organized into multiplexed transcriptional units resembling CRISPR arrays, and often synergize with co-encoded anti-CRISPR proteins to disable hierarchical host immune responses. Furthermore, we demonstrate that these compact viral crlRNA architectures can be harnessed for efficient mammalian genome editing. Our findings uncover widespread viral hijacking of host Cas proteins via crlRNAs, highlighting the recurrent exaptation of CRISPR-Cas regulatory functions to counter multilayered bacterial immune defenses.

## INTRODUCTION

In prokaryotes, CRISPR-Cas systems provide adaptive immunity against mobile genetic elements (MGEs), such as viruses and plasmids^1–3^. These systems fall into two classes, encompassing 7 types (types I-VII) and over 40 subtypes^4–6^. CRISPR arrays consist of unique spacer sequences flanked by direct repeats. Spacers are segments of MGE DNA that are captured and inserted between repeats by a dedicated Cas protein complex, the adaptation module of CRISPR-Cas systems. The CRISPR arrays are transcribed as long precursor RNAs that are processed into mature CRISPR (cr) RNAs consisting of a single spacer flanked by repeat-derived remnants. Mature crRNAs guide Cas effectors to recognize and degrade the reinfecting MGE DNA, which depends on crRNA-protospacer complementarity and the presence of a protospacer-adjacent motif (PAM)^7,8^.

Beyond defense, Cas effectors also regulate host gene transcription via crRNA-like RNAs (crlRNAs) encoded in mini-arrays, which guide Cas proteins to target promoters^9–12^. Unlike crRNAs, crlRNAs enable Cas proteins to bind but not cleave target DNA due to partial complementarity to the target sequences. We previously identified regulatory crlRNAs, termed Cas-regulating (CreR) RNAs, which are typically encoded upstream of type I or V-A *cas* genes and direct Cas effectors to auto-repress their promoters (Figure 1A)^9^. This regulatory mechanism mitigates the risk of CRISPR autoimmunity under normal conditions but allows rapid Cas production when viral anti-CRISPR (Acr) proteins inactivate host Cas proteins^9^. Similarly, long-form trans-activating crRNAs (tracr-L) in type II-A CRISR-Cas systems function as regulatory crlRNAs that mediate Cas9 autoregulation and boost Cas9 expression in response to Acrs^13,14^.

**Figure 1.**
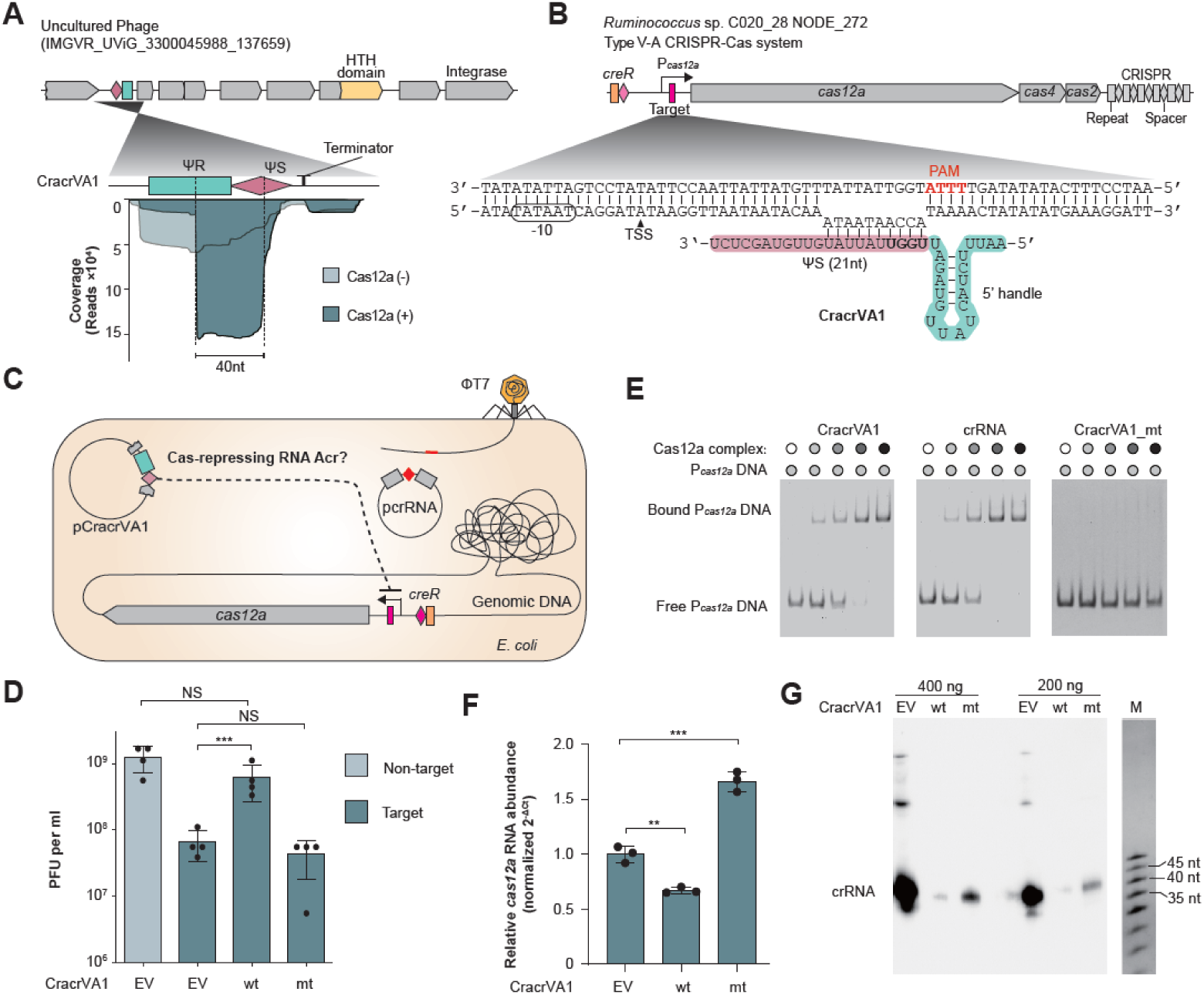
CracrVA1 RNA inhibits transcription of *cas12a*. (A) Schematic of the viral Cracr gene and small RNA-seq analysis of its RNA products. The Cracr gene was introduced into *E. coli* cells expressing or lacking codon-optimized *Ruminococcus* sp. C020_28 Cas12a, and small RNA were extracted for deep sequencing. (B) Complementarity between Cracr RNA and the *cas12a* promoter from *Ruminococcus* sp. C020_28. PAM, protospacer adjacent motif. TSS, transcription start site of *cas12a* (see Figure S1A). Four PAM-proximal nucleotides of the Cracr spacer (bold) were subjected to mutational analysis. (C) Experimental design for assessing the anti-CRISPR effect of CracrVA1. The native CRISPR-Cas locus of BW25113 was replaced with the type V-A locus from *Ruminococcus* sp. C020_28. The pcrRNA plasmid carries either a T7 phage-targeting or non-targeting V-A mini-CRISPR. (D) Plaque forming units of T7 infecting cells expressing wild-type (wt) or spacer-mutated (mt) CracrVA1. EV, empty vector. (E) EMSA of Cas12a co-expressed with different RNA guides. FAM-labelled *cas12a* promoter DNA was used as the probe. (F) Relative *cas12a* mRNA levels (normalized to *ssrA*) determined by quantitative RT-PCR. Error bars, mean±s.d. (n=3). *P* values from two-sided Student’s *t* test. **, *P*≤0.01; ***, *P*≤0.001; NS, not significant. (G) Northern blot detection of phage-targeting crRNAs in Cas12a-bound RNA fractions, with 400 ng or 200 ng of total immunoprecipitated RNA loaded per lane. M, ssRNA marker. See also Figure S1 and S2

Some type I-B or I-F CRISPR-Cas systems encode CreR-like antitoxins (CreA RNAs), which form complexes with the effector Cas proteins and transcriptionally inhibit a toxin (CreT) encoded adjacent to Cas effectors (Figure 1A)^11,15,16^. These CRISPR-regulated toxin-antitoxin (CreTA) modules induce dormancy or cell death when Cas function is compromised by Acrs^15^. Indeed, we have shown that CreTA systems safeguard CRISPR-Cas systems and induce population-level immunity upon sensing Acrs, as a result, providing a broad-spectrum anti-anti-CRISPR function^15^. This strategy is reminiscent of the immune guard hypothesis described in higher eukaryotes and increasingly recognized in prokaryotes, where the inhibition of one defense mechanism triggers the activation of another^17–19^. However, the mechanisms by which viruses evade these hierarchical, multilayered immune defenses remain poorly understood.

Our recent bioinformatic study identified virus-derived small RNAs that resemble bacterial crlRNAs, suggesting a role in CRISPR-Cas repression^12^. However, their mechanisms and biological functions remained unclear. Here, we experimentally demonstrate that diverse viral crlRNAs hijack host type I or subtype V-A Cas proteins to suppress Cas expression or CreT toxin production. To distinguish these viral elements from cellular crlRNAs, we term them Cracrs (<u>C</u>as/CreT-repressing <u>R</u>NA <u>a</u>nti-<u>CR</u>ISPRs), along the same lines as Racrs (RNA anti-CRISPRs), which sequester Cas proteins in aberrant complexes^20^. Mechanistically, Cracrs not only compete with immune crRNAs for Cas binding but also reprogram Cas proteins to transcriptionally repress their own expression or CreT toxin production. Remarkably, Cracr RNAs are frequently arranged in CRISPR-like arrays, enabling viruses to inhibit multiple CRISPR-Cas variants or CreTAs simultaneously. Furthermore, we uncover the functional synergy between type I-F Cracr RNAs and co-encoded AcrIF24, a protein inhibitor that elegantly differentiates Cracr RNAs from host crRNAs/crlRNAs based on their shorter spacer sequences. Our findings uncover a distinct layer of viral counterdefense, showing how viruses combine RNA- and protein-based strategies to evade the multi-layered host immunity. We further show that the compact viral crlRNA architectures can be readily reprogrammed for efficient bacterial and mammalian genome editing applications. Therefore, this work not only expands our understanding of viral immune evasion, but also suggests directions for improving CRISPR-Cas technologies and the design of CRISPR-resistant therapeutic phages against pathogenic bacteria.

## RESULTS

### A viral Cracr RNA inhibits V-A CRISPR-Cas immunity

During the analysis of virus-encoded type V mini-CRISPRs, we identified, in an uncultured phage, a degenerated crlRNA element consisting of a single CRISPR repeat-like (ψR) sequence, a spacer-like (ψS) sequence and a putative rho-independent terminator (Figure 1A). The ψS in this element contains 10 consecutive nucleotides matching the Cas12a promoter (P*_cas12a_*) sequence of *Ruminococcus* sp. C020_28 NODE_272 and the target site is flanked by a type V-A PAM (5′-TTTA-3′) (Figure 1B and Figure S1A), indicating a potential for binding. Notably, upstream of P*_cas12a_* is a putative CreR element that also contains a single ψR, a rho-independent terminator, and a ‘spacer’ sequence with significant similarity to the viral crlRNA (Figure 1B and Figure S1B). The 20 5′-terminal nucleotides of the bacterial and viral ψR sequences are identical to the corresponding portion of the CRISPR repeat from the same locus, which are essential for the formation of the conserved hairpin structure required for Cas12a binding and crRNA maturation (Figure S1C-D). Based on our previous findings on the CreR-directed autoregulation of CRISPR effector protein(s)^9^, we hypothesized that this phage-derived crlRNA hijacks the host regulatory circuit to repress Cas12a expression, and thus designate it CracrVA1 (Cracr for “<u>C</u>as-repressing RNA <u>a</u>nti-<u>CR</u>ISPR”).

We introduced the viral *cracrVA1* gene under the control of its native promoter into *E. coli* cells expressing *Ruminococcus* Cas12a, and confirmed Cas12a-dependent processing of the CracrVA1 precursor RNA by small RNA sequencing (sRNA-seq) (Figure 1A). The 40-nt mature CracrVA1 consists of a 19-nt hairpin-forming 5′ handle followed by a 21-nt spacer (guide) (Figure 1B), similar in size to the cognate mature bacterial CreR RNA (42-nt) (Figure S1E). To assess the predicted anti-CRISPR activity of CracrVA1, we integrated *cas12a* and its native promoter (including the CreR element) into the *E. coli* chromosome and engineered a mini-CRISPR targeting coliphage T7 (Figure 1C). As expected, in the absence of CracrVA1, expression of the phage-targeting crRNA markedly reduced T7 phage forming units (PFUs) compared to a non-targeting control (Figure 1D). CracrVA1 expression restored phage infectivity to near non-targeting levels, demonstrating robust anti-CRISPR activity. Mutation of the four PAM-proximal nucleotides of the Cracr spacer abolished this activity (Figure 1D), underscoring the importance of the spacer identity.

### CrarcVA1 reprograms Cas12a to repress its own transcription

To probe the inhibitory mechanism of CracrVA1, we tested whether it could guide Cas12a to bind dsDNA targets. Electrophoretic mobility shift assays (EMSA) showed that CracrVA1 RNA directed Cas12a to bind its own promoter (P*_cas12a_*) DNA, with an affinity comparable to Cas12a complexed with a typical V-A crRNA designed to bind P*_cas12a_* (Figure 1E). Notably, however, a target DNA cleavage assay revealed that only the crRNA, not CracrVA1, enabled Cas12a to cleave the target DNA (Figure S2A). As expected, Cas12a complexed with a spacer-mutated version of CracrVA1 failed to bind P*_cas12a_* DNA (Figure 1E). This indicates that CracrVA1 reprograms Cas12a to autorepress its own expression in a sequence-specific manner, similar to the previously reported CreR RNAs^9^. We next quantified this repression by quantitative PCR. Compared to an empty vector control, expression of CracrVA1 under its native promoter caused a modest but significant decrease (∼32%, *P* =0.0020) in *cas12a* transcript levels (Figure 1F). Intriguingly, however, expression of the spacer-mutated CracrVA1 resulted in a substantial increase in *cas12a* transcription (by ∼66% compared to the vector control, *P* = 0.0006). We previously showed that CreA-guided suppression of CreT toxins can be similarly relieved by Racrs that sequester Cas proteins^15^. We therefore propose that wild-type CracrVA1 mimics CreR, redirecting Cas12a to enhance autorepression, whereas the spacer mutant relieves autorepression by sequestering Cas12a in non-targeting complexes. Consistent with this model, Western blotting showed that wild-type CracrVA1, but not the spacer mutant, markedly reduced Cas12a protein levels (Figure S2B).

The modest decrease in Cas12a transcription contrasted sharply with the nearly complete loss of phage resistance. We therefore hypothesized that CracrVA1 acts not only by enhancing Cas12a autorepression but also by titrating Cas12a away from immune crRNA complexes toward regulatory ones. To test this hypothesis, we immunoprecipitated cellular Cas12a complexes and analyzed co-purified RNAs by Northern blotting. As expected, expression of wild-type CracrVA1 dramatically reduced the proportion of immune crRNAs among the Cas12a-associated RNAs (Figure 1G), likely, accounting for the observed near complete loss of phage resistance (Figure 1D). The spacer-mutated CracrVA1 also caused reduction of Cas12a-bound crRNA, but to a much lesser extent, the crRNA occupancy of Cas12a being 5-10-fold greater than that observed with the wild-type CracrVA1 (Figure 1G). This finding validates the competition between the viral Cracr RNAs and crRNAs, and further suggests that derepression of Cas12a by the mutated CracrVA1 increased the pool of Cas12a available to crRNAs, consistent with the restored immunity upon CracrVA1 mutation. Together, these findings demonstrate that CracrVA1 antagonizes type V-A CRISPR-Cas immunity by reprogramming Cas12a for autorepression, reducing Cas12a abundance, while simultaneously outcompeting crRNAs for Cas12a binding.

### Systematic discovery of viral Cracr RNAs

Next, we conducted a systematic search for viral crlRNAs across the broader diversity of CRISPR-Cas systems. Our previous study demonstrated the widespread of bacterial crlRNAs, CreR, and CreA, among type V-A and type I CRISPR-Cas loci^9^. Using this CreA/R RNA collection (Table S1) as queries, we performed BLASTn searches in the NCBI RefSeq and IMGVR databases. After excluding hits located within CRISPR-Cas loci (likely CreA/R) and those lacking sufficiently extensive nucleotide matching (at least 9 PAM-proximal base pairs) with the spacer portion of query CreA/R sequences (likely Racrs), we compiled a list of 219 candidate *cracr* genes (each originating from a distinct genome). Among these putative Cracrs, ∼90% (202 out of 219) were from the *Caudoviricetes* class of viruses, with 16 coming from plasmids and one from a phage of *Megaviricetes* (Table S2 and Figure 2A). Based on their correspondence to CreA/R (the BLAST queries), these Cracr candidates were predicted to target type V-A, I-F, I-E, I-C, and I-B CRISPR-Cas loci from a broad taxonomic range of bacteria, spanning 11 orders across the phyla *Pseudomonadota*, *Bacillota*, and *Actinomycetota* (Figure 2A), suggestive of an important role of these viral anti-CRISPR RNAs in the ongoing arms race between diverse bacteria and their genetic parasites.

**Figure 2.**
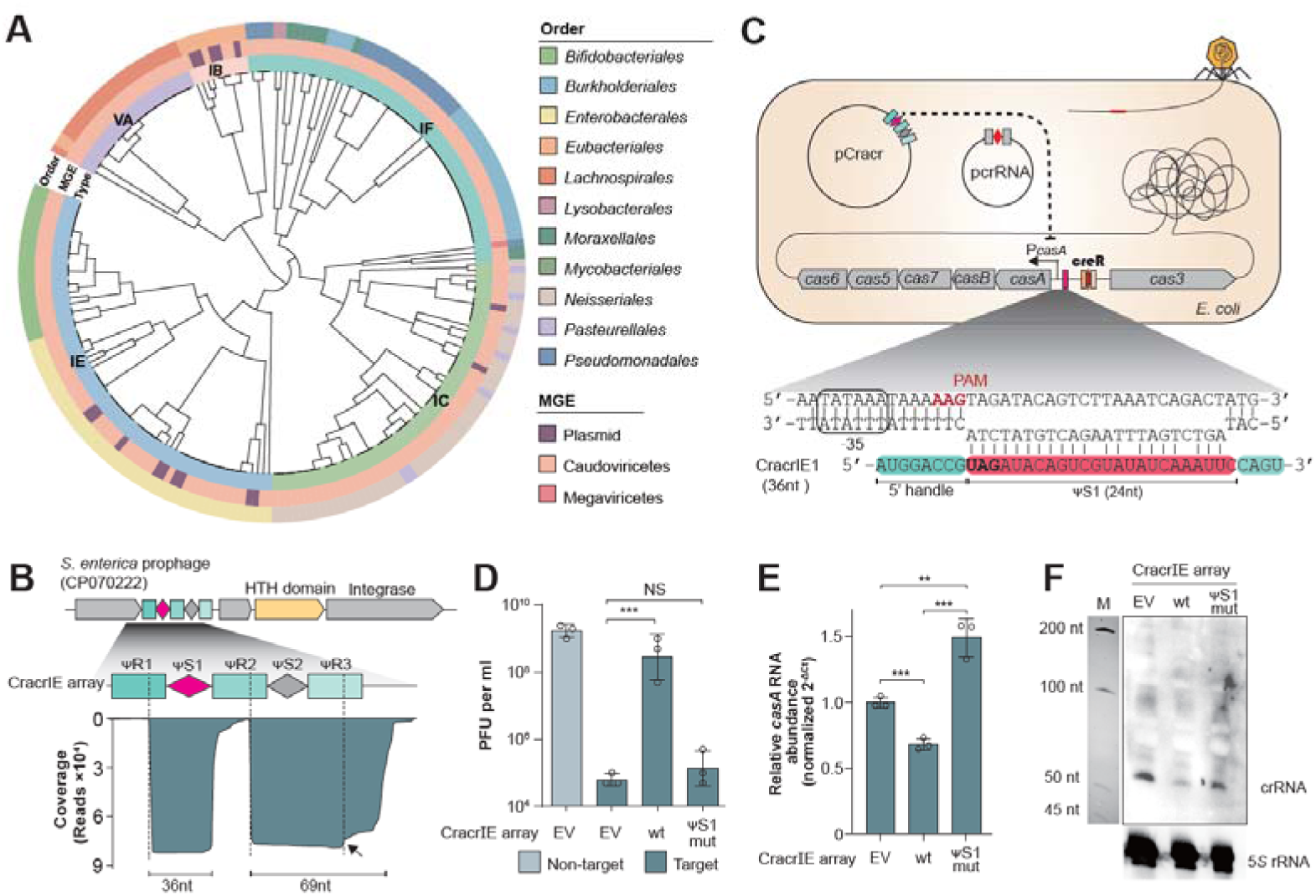
Systematic discovery of viral Cracr genes and a dual-spacer candidate transcriptionally silencing I-E CRISPR-Cas. (A) Clustering of identified Cracr RNAs based on the similarity of their ΨR sequences. The inner ring indicates the origination of each Cracr from different MGEs, while the outer ring depicts the taxonomic distribution of their target CRISPR-Cas loci. (B) Schematic of the prophage-born type I-E Cracr array and small RNA-seq analysis of its RNA products. The array was introduced into *E. coli* BW25113 Δ*cas*Δ*hns* cells expressing the I-E Cascade from *S. enterica* FDAARGOS_717, and small RNA was extracted for deep sequencing. (C) Experimental design for assessing the anti-CRISPR function of the CracrIE array. The native CRISPR-Cas locus of *E. coli* BW25113 was replaced with the *S. enterica* FDAARGOS_717 locus that is targeted by Cracr1. The pcrRNA plasmid carries either a T7 phage-targeting or non-targeting I-E mini-CRISPR. (D) Plaque forming units of T7 infecting cells expressing the wild-type (wt) I-E Cracr array or its ΨS1-mutated variant (altered nucleotides in bold in **C**). EV, empty vector. (E) Relative *casA* mRNA levels (normalized to *ssrA*) determined by quantitative RT-PCR. Error bars, mean±s.d. (n=3). *P* values from two-sided Student’s *t* test. **, *P*≤0.01; ***, *P*≤0.001; NS, not significant. (F) Northern blot detection of phage-targeting crRNAs in cellular total RNAs. M, ssRNA maker. See also Figure S3-S5

We found that 64 of the 219 Cracrs were encoded within 30 arrays containing at least two different spacer sequences (Table S2). This stacked configuration resembles a multi-spacer CRISPR array; henceforth, we refer to these as ‘Cracr arrays’. For most of the arrayed Cracr RNA species, putative target CRISPR-Cas loci could be readily identified based on the sequence similarity to the CreR/CreA queries used in our BLAST search. Examination of contexts of these Cracr arrays within the respective viral genomes suggested that some of them could be parts of larger arrays, carrying additional spacers for which the target CRISPR-Cas loci remained difficult to predict. For example, one metagenomic virus genome was found to carry a three-spacer array in which the first two spacers are predicted to target the I-F CRISPR-Cas systems of *Acinetobacter baumannii* SIMBA061 and W068, respectively, whereas the target of the third remains unidentified (Figure S3A). Notably, Cracr RNAs co-encoded within a single array always target different CRISPR-Cas variants of the same type (usually from closely related strains). For example, one viral array encodes two Cracr RNAs that target two distinct type I-C CRISPR-Cas loci from two *Neisseria sicca* strains (Figure S3B). Altogether, 30 Cracr arrays target I-C (9 two-spacer arrays), I-E (16 two-spacer and 4 three-spacer arrays), or I-F (1 three-spacer array; Figure S3A) CRISPR-Cas systems (Table S2). Thus, viral regulatory crlRNAs have the potential for multiplexing.

### Phage Cracr array silences *Salmonella* I-E CRISPR-Cas

We synthesized a prophage-born *cracr* array from *S. enterica* subsp. *enterica* strain Se29 (Figure 2B), which contains two spacer-like sequences, ψS1 and ψS2, each matching one of the distinct CreR elements embedded within the type I-E CRISPR-Cas loci from *S. enterica* strains FDAARGOS_717 and CFSAN001011, respectively (Table S1 and Figure S4A). Following the introduction of this array into *E. coli* cells expressing Cas proteins from *S. enterica* FDAARGOS_717, sRNA-seq analysis confirmed Cas-dependent maturation of two distinct I-E Cracr RNAs, primarily, 36 nt and 69 nt in length, respectively (Figure 2B). The 36-nt Cracr, hereafter CracrIE1, consisted of a typical 8-nt 5′ handle, the 24-nt ψS1, and four nucleotides derived from the second repeat-like sequence (ψR2) (Figure 2C), suggesting extensive 3′ trimming. In a sharp contrast, the second Cracr, hereafter CracrIE2, contained the 5′ handle, the 21-nt ψS2, and an extended 40-nt 3′ tail. Notably, the third repeat-like sequence (ψR3) harbored a single-nucleotide deletion relative to the canonical CRISPR repeats and other ψR sequences in the array (Figure S4B). This deletion likely resulted in a smaller loop within the RNA hairpin structure recognized by the Cas6 endonuclease (Figure S4C) and hence caused inefficient processing of this repeat unit, as evidenced by the sRNA-seq data (Figure 2B).

Because the fully processed CracrIE1 showed a discontinuous but extensive (20 bp) match to the *S. enterica* FDAARGOS_717 *casA* promoter (P*_casA_*) (Figure S5), the target of its cognate CreR^9^, we integrated the Cascade- and Cas3-encoding genes of this strain into the *E. coli* chromosome and engineered a type I-E mini-CRISPR targeting T7 to assess anti-CRISPR activity of this Cracr (Figure 2C). In the absence of the viral Cracr array, expression of the phage-targeting crRNA reduced T7 PFUs by approximately 4 logs compared to a non-targeting control (Figure 2D). By contrast, co-expression of the CracrIE array almost fully restored phage infectivity. Mutating ψS1 abolished this effect (Figure 2D), confirming that complementarity of CracrIE1 to P*_casA_* is essential for its inhibitory function.

Quantitative PCR showed that the wild-type CracrIE array reduced *casA* transcription by about 30% compared to the empty vector control, whereas the ψS1-mutated CracrIE array, conversely, caused an about 50% increase in transcription (Figure 2E), recapitulating the pattern observed for CracrVA1 (Figure 1F). Northern blotting further showed that the proportion of phage-targeting crRNAs in the total cellular RNAs was substantially reduced upon introduction of the wild-type CracrIE array, but not the ψS1-mutated version (Figure 2F). We propose that, analogous to CracrVA1, the CracrIE array not only reprograms the Cas effector to tighten its autorepression, but also produces RNAs (likely, both CracrIE1 and CracrIE2) that compete with crRNAs for Cas proteins. Although aberrant processing of the ψS2-derived Cracr species precluded functional validation, its predicted targeting of a distinct *Salmonella* type I-E locus points to the multiplexing potential of Cracr arrays and suggests that phages deploy them to counter diverse CRISPR systems in their hosts.

### Widespread association of *acrIF24* with I-F Cracrs

We noticed that 21 of the 58 I-F CRISPR-Cas-targeting *cracr* genes are tightly associated with the *acrIF24* gene encoding a known I-F-specific Acr protein^21^ (Figure 3A and Table S2), suggesting that *acrIF24* could serve as a marker for discovering additional I-F-targeting Cracr elements. We therefore performed genomic context-based searches using *acrIF24* as a bait and searched for Cracr within the flanking regions. This approach identified 221 candidate *cracr* genes, of which only three were identified in our initial screening based on similarity to CreR/CreA (Table S3), highlighting the underestimated abundance of these elements. Remarkably, most *acrIF24* genes are accompanied by a candidate *cracr* gene, and this genetic linkage shows a broad taxonomic distribution, spanning 8 bacterial genera from 6 orders of β- and γ-proteobacteria (Figure 3B).

**Figure 3.**
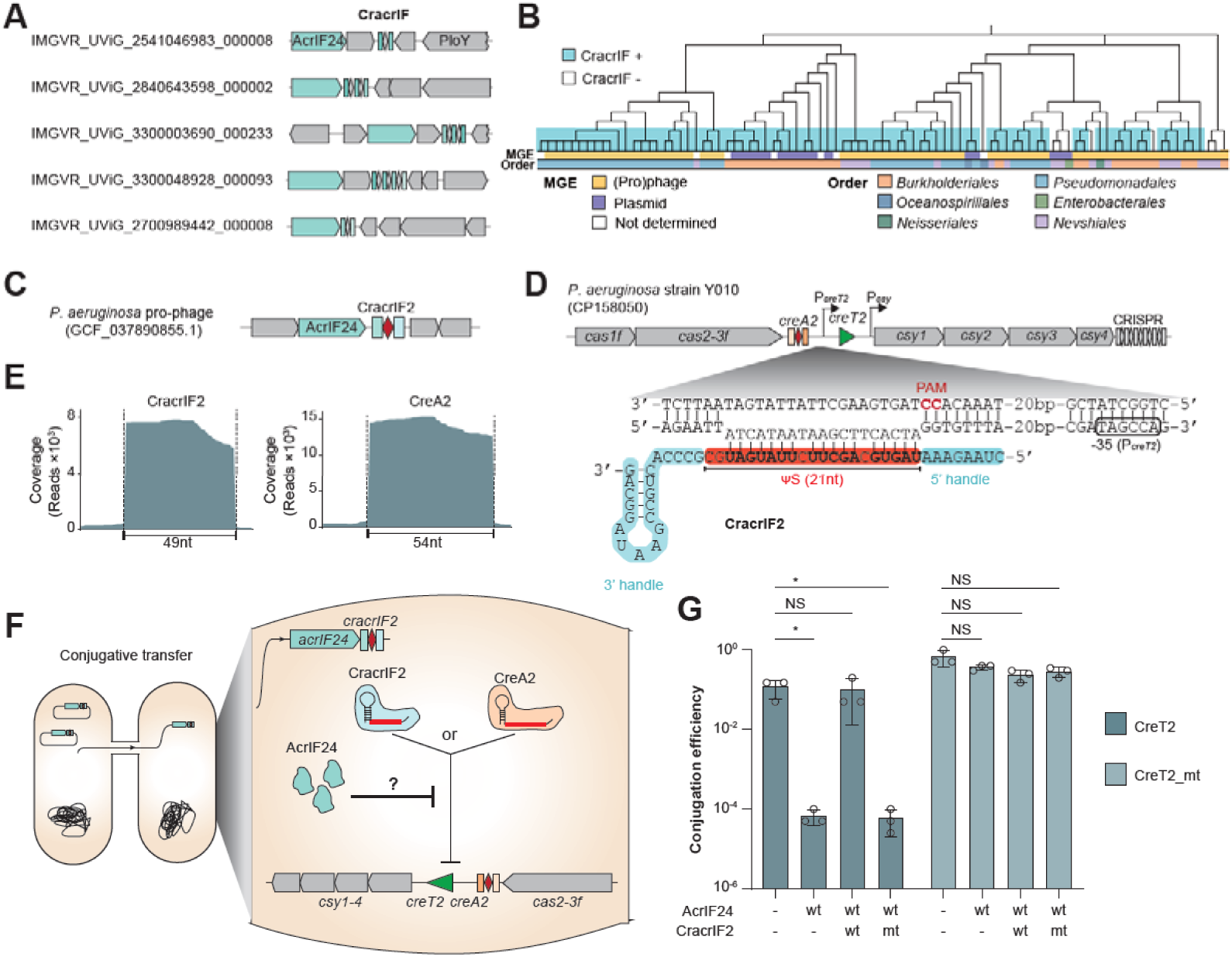
CracrIF suppresses CreTA-based immunity. (A) Five representative *acrIF24*-associated CracrIF genes/arrays carried by MGEs from diverse bacterial species. (B) Genetic association of AcrIF24 proteins to CracrIF RNAs across different phylogenetic orders. Tree is constructed from all non-redundant AcrIF24 proteins in NCBI RefSeq, IMG/VR, and IMG/PR databases. (C) Schematic of the *P. aeruginosa* prophage-encoded *cracrIF2* and associated *acrIF24*. (D) *P. aeruginosa* I-F CRISPR-Cas locus harboring the CreTA module (named CreTA2^22^) targeted by CracrIF2. Transcription start sites (TSSs) of *creT2* and *csy* operon were mapped in Figure S7B. (E) Small RNA-seq revealing different sizes of CracIF2 and CreA2. (F) Schematic of the conjugation assay design. The recipient strain is a modified PA14 derivative harboring the Y010 CreTA2 module integrated into its chromosomal CRISPR-Cas locus. (G) Conjugation efficiency of a plasmid encoding AcrIF24 alone or together with either wild-type (wt) or spacer-mutated (mt) CracrIF2. The recipient strain encodes either wild-type CreT2 or its proline-codon mutant (CreT2_mt). Error bars, mean±s.d. (n=3). *P* values from two-sided Student’s *t* test. *, *P*≤0.05; NS, not significant. See also Figure S6-S8

The *cracr* genes identified through the association with *acrIF24* show a higher proportion of multi-spacer arrangements (76 of the 221 contained more than one spacer), collectively encoding 367 putative Cracr RNAs (including sequence-redundant variants) (Table S3). Although some of these Cracr RNAs share similar or even identical spacer sequences, their ΨS sequences are diverse, suggesting the ability to target a range of CRISPR-Cas loci. Indeed, spacer-protospacer matching analysis allowed us to predict targets consistently mapping to *cas* promoters within CRISPR-Cas loci for 96.55% (140 of the 145) of single-spacer *cracr* genes and for at least one spacer in 69.74% (53 of the 76) of multi-spacer *cracr* arrays (Table S3). Notably, among the spacers with confidently predicted targets, those within the same Cracr array consistently targeted CRISPR-Cas loci of the same subtype. Together, these findings suggest that Cracr RNAs are widely employed by phages and plasmids to counteract type I-F CRISPR-Cas systems.

### A CracrIF RNA suppresses CreTA-based population-level immunity

The genetic linkage between AcrIF24 and Cracr RNAs targeting type I-F CRISPR loci suggests the possibility of functional synergy between protein- and RNA-based anti-CRISPRs. To explore this possibility, we focused on anti-CRISPR elements carried by *P. aeruginosa* MGEs. We recently demonstrated that I-F CRISPR-Cas loci in *P. aeruginosa* are frequently (∼26%) associated with either one of two crlRNAs, CreA1 and CreA2, which inhibit the promoter of conserved proline-codon-containing RNA toxins that safeguard CRISPR-Cas immunity (CreT1 and CreT2, respectively)^22^ (Figure S6A). The spacers of all identified Cracr RNAs encoded by *P. aeruginosa* MGEs are notably conserved (Table S2 and 3), and share high sequence similarity with either CreA1 or CreA2 (examples shown in Figure S6B-C; hereafter, CracrIF1 and CracrIF2, respectively), suggesting that these viral RNAs silence the two CreTA elements by mimicking their regulatory crlRNA antitoxin components. We synthesized two of these *cracr* genes (Figure S6B-C) and introduced them into *P. aeruginosa* PAO1 cells for characterization. Northern blot analysis confirmed maturation of CracrIF1 and CracrIF2, which was dependent on Csy1-4, and demonstrated their smaller sizes (46 and 49 nt, respectively) compared to cognate CreA RNAs (48 and 54 nt, respectively) (Figure S6D-E). Subsequent functional interrogation using an mCherry-based reporter assay showed that CracrIF1 and CracrIF2 strongly inhibited transcription from their cognate target promoters (P*_creT1_* and P*_creT2_*, respectively) in PAO1 cells (Figure S6F).

We then sought to investigate whether CracrIF elements inhibited the CRISPR-safeguarding activity of CreTA systems in *P. aeruginosa*. To avoid polar effects arising from shared promoters with adjacent *cas* genes in the CreTA1 locus (Figure S7), we focused on the independently expressed CreTA2 system associated with the type I-F CRISPR-Cas locus from strain Y010 (Figure 3D), which is targeted by CracrIF2. We first introduced the Y010 CRISPR-Cas locus (encoding CreTA2) into PAO1 cells and transformed these cells with a plasmid expressing AcrIF24, either with or without co-expression of CracrIF2 (Figure S8A). In the absence of CracrIF2, induced expression of AcrIF24, which inhibits the target DNA-binding activity of I-F Cascade^21^, resulted in a marked reduction (>4-log) in colony forming unit (CFU) relative to the empty vector control (Figure S8B), suggesting that AcrIF24 triggers CreT2 expression, thereby impairing host cell growth. Consistently, mutation of the two critical proline codons in *creT2* fully restored colony growth. Co-expression of CracrIF2 with AcrIF24 also restored the CFU to control levels, and this effect was strictly dependent on complementarity between CracrIF2 and P*_creT2_* (Figure S8B), consistent with inhibition of CreT2 expression by CracrIF2. Small RNA-seq further confirmed the production of the 49-nt CracrIF2, as well as the 54-nt CreA2 RNAs (Figure 3E).

We next conducted conjugation assays to examine whether AcrIF24 and CracrIF2 act during the initial delivery of incoming DNA (Figure 3F). As expected, a conjugative plasmid carrying *acrIF24* was poorly transferred into a recipient harboring CreTA2, but efficiently into a modified recipient with mutated *creT2* (Figure 3G), indicating that AcrIF24 triggers CreTA2-driven growth arrest upon entry. Co-delivery of CracrIF2, but not its mutant, abolished this conjugation block, consistent with the CFU recovery data from inducible expression assays (Figure S8B). Thus, AcrIF24 and CracrIF2 function as a tightly coordinated anti-defense module that rapidly disarms CreTA2-mediated immunity upon entering the bacterial cell. Specifically, while AcrIF24-mediated Cascade inhibition relieves CreA-guided repression of *creT* transcription, triggering toxicity and blocking gene transfer (or phage replication, in the native system), co-expressed CracrIF2 targets the *creT* promoter, rescuing the cells and suppressing the side-effect of AcrIF24, highlighting the coordination between RNA- and protein-based anti-CRISPR elements. More broadly, this exemplifies how phages overcome hierarchical host immune responses by deploying multilayered, synergistic counter-defense.

### AcrIF24 differentiates between CreA2 and CracrIF2 by spacer size

We reasoned that AcrIF24 might be able to discriminate between Cas complexes assembled with viral Cracrs and those assembled with bacterial crlRNAs through an unknown mechanism that enables Acr-Cracr synergy. Indeed, using a P*_creT2_*-driven mCherry reporter assay, we found that AcrIF24 almost completely abolished transcription repression by CreA2-guided Cas complexes, whereas CracrIF2-guided silencing remained unaffected (Figure S9A). Comparing the sequences of *cracrIF2* and *creA2* genes, we hypothesized that AcrIF24 discrimination relies on their different ΨR sequences, ΨS sizes, or a combination of both (Figure 4B). To test this prediction, we replaced the ΨR sequences of *cracrIF2* with those of *creA2* (Figure S9B). However, the mCherry assay showed that the mutant remained resistant to AcrIF24, ruling out ΨR sequence differences as a key determinant (Figure S9B). Next, we extended the ΨS of *cracrIF2* from 21 nt to 26 nt, to match *creA2*, and conversely, truncated the ΨS of *creA2* to match *cracrIF2*. The Csy complexes guided by the ΨS-extended *cracrIF2* became highly susceptible to AcrIF24 inhibition, whereas the ΨS-truncated *creA2* became resistant (Figure 4B), indicating that ΨS size is mechanistically critical. Indeed, a survey of diverse type I-F *cracr* genes (n = 76) and their bacterial counterparts (*creA* or *creR*) (Table S3) revealed a significant difference in ΨS sizes (*P* = 6.23E-08) (Figure 4C). Viral Cracr RNAs contain ΨS sequences primarily between 17-21 nt in size, whereas bacterial CreA/CreR RNAs typically range within 23-27 nt. To further investigate the role of ΨS size in AcrIF24 discrimination, we generated CracrIF2 derivatives with ΨS sizes ranging from 18 nt to 26 nt and found that susceptibility to AcrIF24 inhibition markedly increased when the spacer size exceeded 22 nt (Figure S9C). Together, these results reveal an elegant mechanism by which AcrIF24 discriminates between Csy complexes assembled with different crlRNAs based on small differences in spacer length, thereby enabling functional synergy with co-encoded Cracrs.

**Figure 4.**
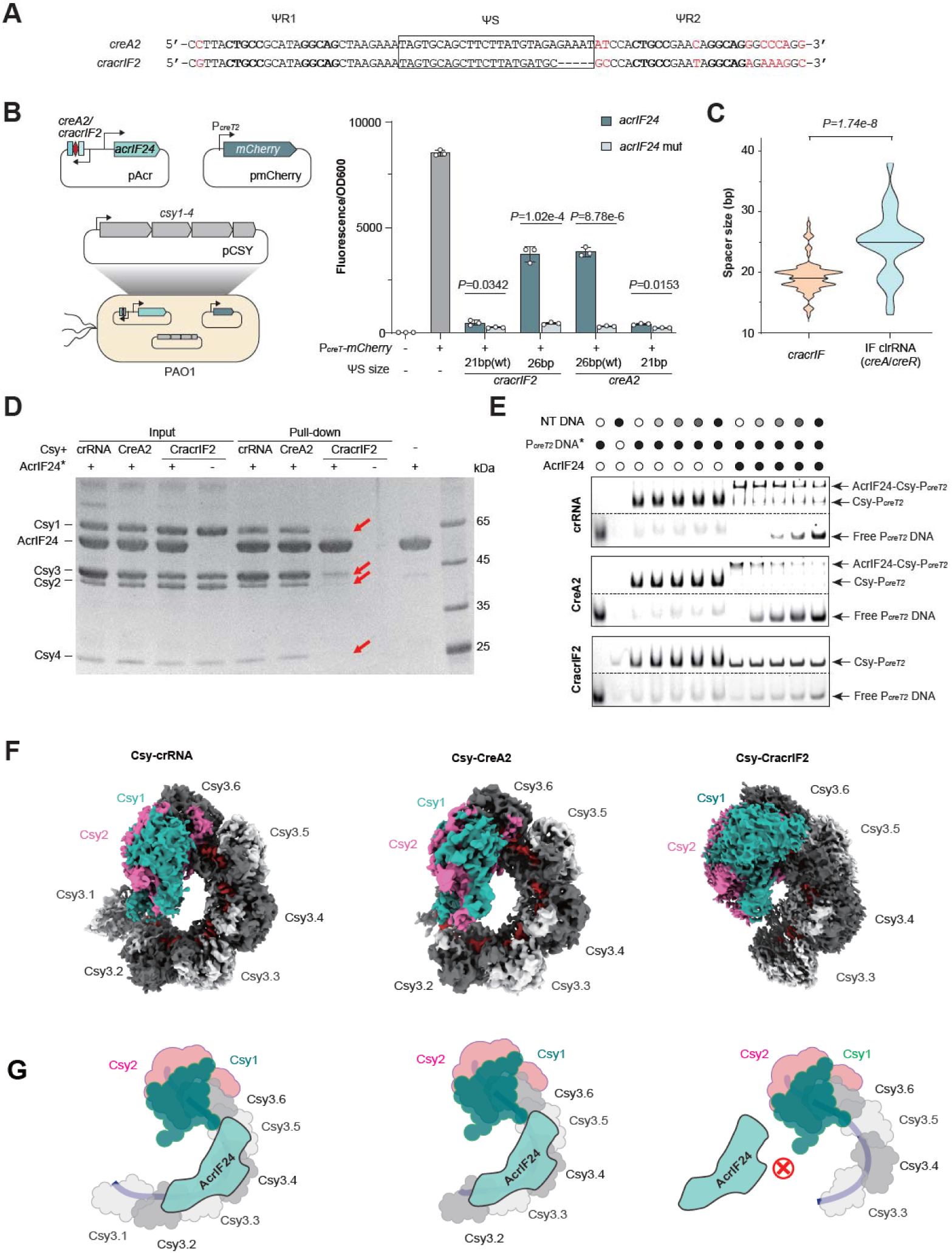
AcrIF24 differentiates between CracrIF2 and CreA2 through their different spacer sizes. (A) Illustration showing the sequence differences between *creA2* and *cracrIF2*. (B) The inhibitory effects of the wt or ΨS-extended (to 26 bp) CracrIF2, or the wt or ΨS-truncated (to 21 bp) CreA2 on P*_creT2_* in the presence of AcrIF24 or its mutant. Error bars, mean±s.d. (n=3). *P* values from two-sided Student’s *t* test. (C) Distribution of the spacer size of all identified *cracrIF* and related *creA/creR* elements. The median size is 19 nt for *cracrIF* and 25 nt for CreR/CreA. The *P* value was determined using the Mann–Whitney *U* test (two-sided). (D) GST-labeled AcrIF24 pull-down experiments with Csy proteins bound to canonical crRNA, CreA2, or Cracr2. The crRNA control is designed based on CreA2 with five extra spacer nucleotides not matching P*_creT2_*, resulting a typical spacer size of 32 nt. Red arrows indicate the inefficiently captured Csy proteins in the CracrIF2 assay. (E) Competition assays with non-target DNA (0, 0.15, 0.3125, 0.625, or 1.25 μM) and FAM-labeled target P*_creT2_* DNA (0.04 μM) for 0.8 μM Csy-crRNA, Csy-CreA2, or Csy-CracrIF2 complexes in the absence or presence of AcrIF24 proteins (9.6 μM). (F) Solved Cryo-EM structures of the Csy complex assembled with crRNA, CreA2, or CracrIF2. The Csy3 subunits are numbered according to the canonical Csy complex with crRNA. (F) Scheme illustrating the ability of AcrIF24 to bind the longer Csy-crRNA and Csy-CreA2 complexes, but not the shorter Csy-CracrIF2 complex. See also Figure S9-S14

### AcrIF24 cannot bind Csy-Cracr complexes with fewer Csy3 subunits

The length of crRNAs determines the number of Csy3 subunits in type I-F Csy complexes^23,24^. In PAO1 cells expressing PA14 Csy proteins, sRNA-seq and Northern blot analyses both confirmed the different lengths of CreA2 (54 nt), CracrIF2 (49 nt), and a canonical crRNA (60 nt) (Figure 3E and Figure S6E). We therefore hypothesized that Csy complexes guided by these RNAs assemble with different Csy3 stoichiometries. To explore this possbility, we purified Csy complexes assembled with each backbone RNA using His_6_-Csy1 affinity purification. Size-exclusion chromatography (SEC) revealed distinct elution profiles: Csy-crRNA complexes eluted first, followed by Csy-CreA2, whereas Csy-CracrIF2 eluted last as a markedly shorter and broader peak (Figure S10A-B), suggesting a reduced and heterogeneous Csy3 incorporation. Consistently, native PAGE analysis of Csy-CracrIF2 showed diffuse bands, supporting heterogeneity in Csy3 composition (Figure S10C). SDS-PAGE further confirmed that Csy-CracrIF2 complexes contained fewer Csy3 subunits than Csy-crRNA and Csy-CreA2 complexes (Figure S10D-E).

A previous study showed that AcrIF24 binds Csy-crRNA complexes through extensive interactions with five of the six Csy3 backbone subunits (except Csy3.1)^21^, leading us to hypothesize that AcrIF24 is unable to bind Csy-CracrIF2 complexes, which contain fewer Csy3 subunits. To test this prediction, we performed pull-down assays using GST-tagged AcrIF24. While Csy-crRNA and Csy-CreA2 complexes were efficiently captured, only minimal amounts of Csy-CracrIF2 were recovered, indicating a substantially lower AcrIF24 affinity for Csy-CracrIF2 (Figure 4D). We next investigated the DNA-binding specificity of the different Csy complexes in the presence or absence of AcrIF24. As described previously^21^, AcrIF24 induced Csy-crRNA and Csy-CreA2 complexes to bind non-target DNA, whereas Csy-CracrIF2 remained target-specific (Figure 4E). This observation is consistent with the inability of AcrIF24 to interfere with CracrIF2-directed transcriptional repression (Figure S9A).

To resolve the structural basis of these interactions, we performed cryo-electron microscopy (Cryo-EM) of Csy complexes assembled with the different RNA guides (Figure S11-13). Csy-crRNA and Csy-CreA2 complexes contained six and five Csy3 subunits, respectively, whereas Csy-CracrIF2 primarily assembled with four subunits (Figure 4F). Comparison with the previously reported Csy-crRNA-AcrIF24 structure^21^ revealed that loss of Csy3.1 and Csy3.2—but not Csy3.1 alone—markedly reduced the AcrIF24 binding interface (Figure S14). These results indicate that the functional compatibility of CracrIF2 (and likely CracrIF1) with AcrIF24 arises from their shorter spacer lengths, which preclude incorporation of two Csy3 subunits required for stable AcrIF24 engagement (Figure 4G). Together, these findings reveal a previously unrecognized co-evolution between RNA- and protein-based anti-CRISPRs, resulting in coordinated inhibition of both adaptive and innate immune responses in bacteria.

### Harnessing V-A Cracrs for genome editing in bacteria and human cells

We next investigated whether Cracr RNAs could direct target DNA cleavage upon minimal engineering, through extension of spacer-target complementarity. We first modified CracrVA1 to fully match P*_cas12a_* and introduced it into the *E. coli* cells containing a chromosomal *cas12a* and its native promoter (Figure S15A). This full-matching CracrVA1 variant substantially reduced transformation efficiency compared to the wild-type Cracr, suggesting that this modified viral RNA no longer mediated gene regulation but instead directed cleavage of the bacterial chromosome. To further explore its genome editing potential, we replaced ψS with a spacer fully matching a non-essential *E. coli* gene and introduced it into cells together with a donor plasmid providing the λ-Red recombination system and two homology arms flanking the target locus (Figure S15B). Remarkably, among 22 randomly selected transformants, the target gene was deleted in-frame with an efficiency of 100% (Figure S15C-D), demonstrating the high precision and reliability of this phage-derived RNA-guided editing strategy in bacteria.

Building upon these results in bacteria, we sought to evaluate whether the scaffold of CracrVA1 could be reprogrammed for mammalian genome editing. We designed a CracrVA1 variant with the eukaryotic transcription terminator T6 and a spacer fully matching the target human gene (*FANCF, DNMT1, EXM1*, or *RUNX1*) (Figure S16A). Given that the natural crRNA guide of *Eubacterium rectale* Cas12a (ErCas12a) shares nearly identical 5’ handle nucleotides with the CracrVA1 RNA (Figure S16B), we selected this commercial human-codon-optimized nuclease for our experiments. We co-transfected HEK293T cells with the targeting CracrVA1 construct and an ErCas12a expression plasmid (Figure S16A). Following enrichment of successfully transfected cell populations, Illumina deep sequencing demonstrated robust introduction of indels into each gene, with the editing efficiency of *FANCF* reaching up to approximately 50% (Figure S16C). Notably, these frequencies were comparable to those achieved with conventional crRNA guides, underscoring that the phage-derived CracrVA1 RNA scaffold functions efficiently, without compromising activity, in the complex chromatin environment of mammalian cells. Collectively, these findings establish that the minimized viral Cracr RNA architecture can be harnessed for precision genome engineering in both prokaryotic and mammalian cells.

## DISCUSSION

Disabling host defense systems is a central strategy employed by MGEs to ensure their propagation^25^. In particular, viruses and plasmids of prokaryotes have evolved diverse mechanisms to inhibit CRISPR-Cas immunity, including over 100 identified Acr protein families^26,27^ and the recently discovered RNA Acrs, Racrs^20^. In response, many CRISPR-Cas systems have evolved multilayered defensive architectures including crlRNA-guided autoregulation (e.g., CreR in type V-A and I-E systems) which stabilizes *cas* gene expression via a negative feedback loop, rendering the defense system resistant to conventional Cas-inhibiting Acrs or Cas-sequestering Racrs^9^ (Figure 5A). Additionally, some systems use crlRNAs (e.g., CreA in type I-F) to direct Cas proteins to transcriptionally control a toxin (CreT) gene, which triggers abortive infection when CRISPR-Cas is compromised by Acrs or Racrs^15^ (Figure 5A). The discovery of this multilayered prokaryotic immunity poses the question how MGEs can effectively evade such a robust, buffered defense.

**Figure 5.**
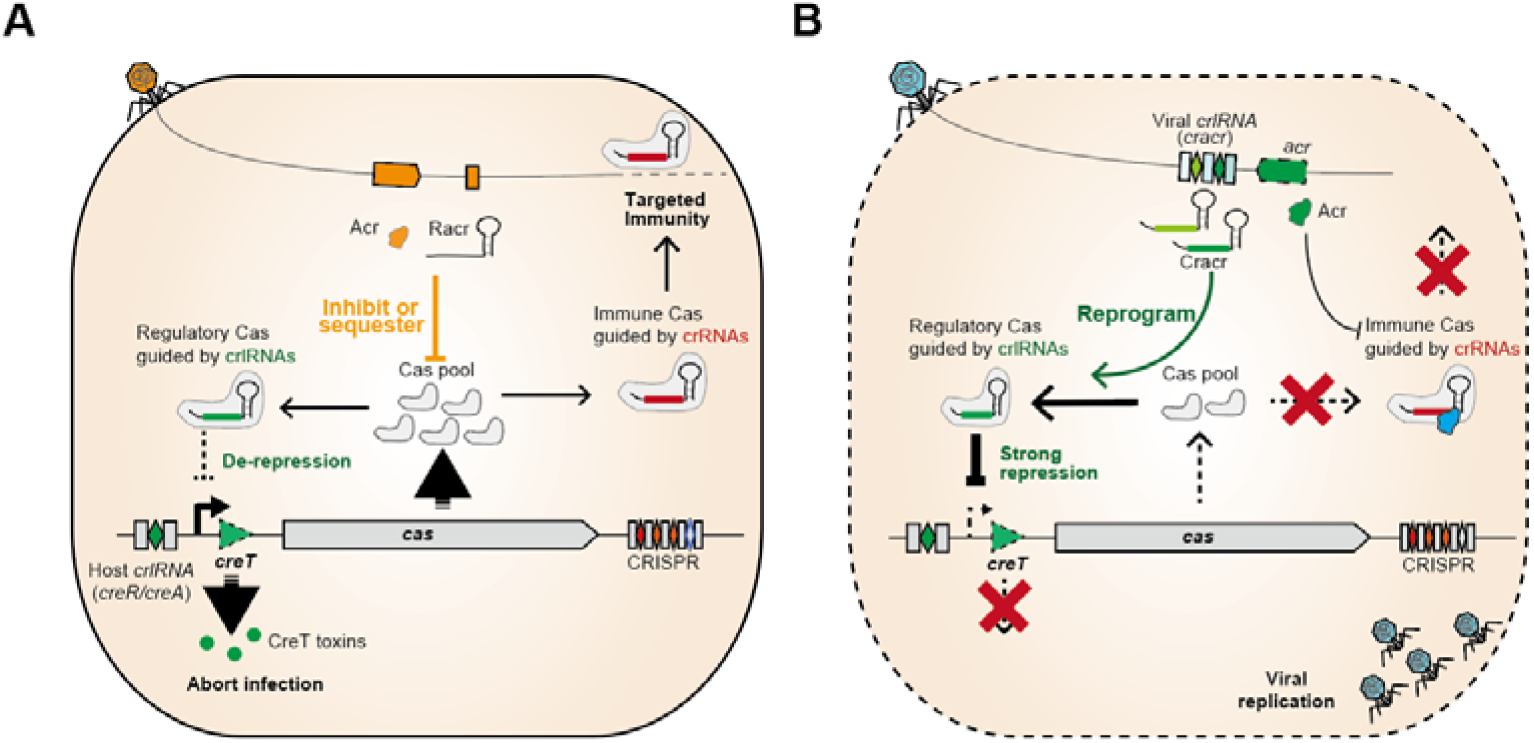
Proposed action modes of viral Cracr RNAs. (A) In type I or V CRISPR-Cas systems equipped with crlRNAs (CreR/CreA) that modulate Cas or CreT expression, viral Acrs or Racrs often fail because Cas inhibition or sequestration leads to de-repression of *cas* or *creT* promoters. This results in a burst of production of Cas proteins or CreT toxins, thereby restoring CRISPR-Cas immunity or triggering abortive infection. (B) Cracr RNAs, viral mimics of CreR/CreA, reprogram Cas proteins to repress *cas* or *creT* promoters, tightening Cas autorepression or CreT repression while simultaneously competing with immune crRNAs for Cas binding. Consequently, these Cas- or CreT-repressing anti-CRISPR RNAs effectively subvert crlRNA-armed CRISPR-Cas systems, particularly, when acting in concert with co-encoded Acrs or Racrs.

Here, we describe a distinct class of viral anti-defensive RNAs, Cracrs (<u>C</u>as/CreT-repressing <u>R</u>NA <u>a</u>nti-<u>CR</u>ISPRs), which counteract bacterial immunity by mimicking host regulatory crlRNAs, thereby reprogramming diverse Cas effectors to silence Cas or CreT promoters. Unlike previously characterized anti-CRISPR elements, such as Acrs that inhibit Cas activity or Racrs that function solely via Cas protein sequestration, Cracrs not only compete with crRNAs for Cas binding but additionally convert the captured Cas effectors into transcriptional repressors that specifically silence Cas or CreT promoters (Figure 5B). This dual capacity allows Cracrs to overcome CreR-armed CRISPR-Cas systems (e.g., type V-A and I-E), where simple sequestration of the effectors would fail due to autoregulatory compensation. Indeed, type V-A and I-E Cracr RNAs redirect Cas effector complexes to repress *cas* gene expression, thereby simultaneously depleting the Cas protein pool and outcompeting immune crRNAs. In systems where bacterial crlRNAs regulate toxin genes (e.g., CreTA in type I-F), a Cracr directly targets the toxin promoter, blocking production of the lethal CreT toxin that would otherwise be derepressed upon Cas effector inhibition. Thus, Cracrs hijack the host’s own regulatory logic to dismantle both layers of the multilayered immune network.

We found that multiple Cracr RNAs often cluster within the same transcriptional unit. This organization of viral Cracr arrays mirrors bacterial CRISPR arrays, highlighting the unique multiplexing potential of CRISPR-Cas systems and the versatility of RNA components leveraged by both prokaryotes and MGEs to orchestrate defense and counter-defense^28^. Cracr arrays can enable MGEs to simultaneously repress different CRISPR-Cas systems in the same or related hosts. Furthermore, this natural multiplexing can enhance the sequestering effect because combined expression of several Cracrs collectively increases the pool of non-immune Cas complexes, thereby more effectively depleting Cas proteins available to form productive complexes with crRNAs.

In addition, we found that CracrIF RNAs synergize with protein Acrs, as suggested by their tight genetic linkage to *acrIF24* (Figure 3). Given that Cracr RNAs guide Cas proteins to bind target promoters, their association with AcrIF24, a potent DNA-binding inhibitor^21^, initially appeared paradoxical. However, we found that Cracr RNAs circumvent this apparent incompatibility by using shorter spacer sequences, resulting in Csy-Cracr complexes lacking two Csy3 subunits required for AcrIF24 binding (Figure 4). Therefore, while AcrIF24 inhibits immune Csy complexes loaded with virus-targeting crRNAs, it does not interfere with regulatory complexes loaded with viral Cracrs, highlighting the co-evolution of RNA- and protein-based anti-CRISPR strategies. As a result, Cracrs can rescue prokaryotic cells from the toxicity of CreT-like (and, likely, other) toxins that are otherwise unleashed when Acrs or Racrs inhibit CRISPR effectors.

Beyond the CRISPR-CreTA coupling, many other cases of multilayered defenses in prokaryotes have been identified, suggesting that this could be a common modality of prokaryotic immunity. For example, Retron Ec48 safeguards RecBCD, a key antiviral complex in bacteria^29,30^, whereas PARIS systems trigger abortive infection in response to phage inhibitors of restriction-modification and BREX systems^31,32^. The frequent clustering of defense systems in bacterial genomes^33–35^ suggests that these systems frequently cooperate to provide hierarchical, multi-layered protection^36^ and recently multiple cases of such synergy have validated experimentally. Collectively, these findings support the proposition that mechanisms involving dormancy or cell death induction comprise the last line of defense in prokaryotes that is activated once active immunity mechanisms fail^37,38^. These discoveries further align with the guard hypothesis initially proposed for immune pathways in higher eukaryotes, where inactivation of one defense triggers the activation of another^17–19^. The findings reported here suggest that, in the course of the perennial arms race, MGEs across domains of life countered the evolution of multilayered immunity in their hosts by evolving multilayered anti-defense strategies.

Complementing our findings on Cracr RNAs targeting type I and V CRISPR-Cas systems, a parallel study uncovered similar regulatory RNA species that silence type II CRISPR-Cas systems^39^. These elements, named viral single-guide RNAs (vsgRNAs), appear to derive from host Cas9-regulating tracr-L RNAs and similarly redirect Cas9 to inhibit its own promoter. Therefore, MGEs have repeatedly hijacked host crlRNAs across CRISPR-Cas diversity. These findings add to the growing body of evidence that both bacteria and diverse MGEs evolved compact RNA-guided transcriptional regulatory functions billions of years before the invention of their human-engineered counterparts^40,41^. Notably, both Cracrs and vsgRNAs can be repurposed for genome editing in bacteria and in human cells.

In summary, our findings reveal that Cracr RNAs represent a distinct class of anti defense elements that function via a dual mechanism: by competitive sequestration of Cas effectors and reprogramming them into transcriptional repressors. This multilayered antidefense strategy can overpower autoregulated or toxin-associated CRISPR Cas systems that are refractory to simple inhibition or sequestration. Furthermore, Cracrs often cooperate with co-encoded protein Acrs, preventing activation of immune toxins upon inhibition of CRISPR effectors. These discoveries expand our understanding of the complexity of viral anti-defense strategies and have implications for developing compact CRISPR-Cas tools for genome editing and CRISPR-resistant therapeutic phages.

## RESOURCE AVAILABILITY

### Lead contact

Further information and requests for resources and reagents should be directed to and will be fulfilled by the Lead Contact, Ming Li.

### Materials availability

Materials are available from Ming Li upon request.

### Data and code availability

This paper does not report original code.

All data and materials reported in this paper will be shared by the lead contact upon request. This paper does not report original code.

## ACKNOWLEDGMENTS

This work was supported by the National Key Research and Development Program of China [2024YFA0919400], the Strategic Priority Research Program of the Chinese Academy of Sciences [XDB0810000], the National Natural Science Foundation of China [32150020, 32230061, 32270092, and 32400063], and the Lundbeck Fonden grant [R347-2020-2346], the research grant from VILLUM FONDEN [VIL60763].

## AUTHOR CONTRIBUTIONS

M.L. conceived and supervised the project with valuable suggestions from R.P.-R and Y.F.. M.L, C.L., Z.L., R.P.-R., Y.F., R.W., and W.W. designed the experiments. C.L., J.Y., J.X., G.L. L.C., M.R.-M., and R.P.-R. performed the bioinformatics analyses and predicted *cracr* genes with valuable suggestions from E.V.K. J.Y., G.L. and C.L. constructed the phylogenetic and clustering trees. C.L. and R.W. generated strains and plasmids. C.L. performed phage infection assays. C.L. and R.W. generated mutants of all strains. C.L. and R.W. performed Northern blot. C.L. J.X and S.X. performed sRNA-seq and 5′ RACE assays. Z.L., X.D., and L.G. conducted protein purification. Y.F. and W.W. resolved the Cryo-EM structure. Z.L. performed *in vitro* EMSA, DNA cleavage, GST pull-down and Western blot assays. F.C. performed RT-qPCR. J.X., J.D. and C.L. tested the editing efficiency of CracrVA1 in human cells and *E. coli*. M.L., C.L., Z.L., R.P.-R, E.V.K. and W.W. analyzed data. M.L., C.L., R.P.-R., Y.F., and Z.L. prepared the original draft. M.L., R.P.-R., Y.F., and E.V.K finalized the manuscript with input from all authors.

## DECLARATION OF INTERESTS

The authors declare no competing interests.

## STAR* METHODS

**KEY RESOURCES TABLE**

Detailed methods are provided in the online version of this paper and include the following:

KEY RESOURCES TABLE

RESOURCE AVAILABILITY

○ Lead contact

○ Materials availability

○ Data and code availability

EXPERIMENTAL MODEL AND STUDY PARTICIPANT DETAILS

○ Bacterial strains and growth conditions

METHOD DETAILS

○ Plasmid construction, transformation, and strain construction

○ Mammalian cell culture, transfection and genomic DNA extraction

○ High throughput sequencing

○ Phage propagation

○ Phage infection assays

○ Dilution plating assay

○ RNA isolation

○ 5′ RACE analysis

○ Small RNA sequencing and analysis

○ RT-qPCR

○ Fluorescence measurement

○ Northern blot

○ Western blot

○ Protein expression and purification

○ Small RNA synthesis

○ DNA cleavage assays

○ Electrophoretic mobility shift assay (EMSA)

○ GST pull-down assay

○ Cryo-EM sample preparation and data acquisition

○ Cryo-EM data processing

○ Cryo-EM model building and refinement

○ Search for *cracr* genes

○ Hierarchical clustering of *cracr* repeat regions

○ Phylogenetic analysis of AcrIF24 proteins

QUANTIFICATION AND STATISTICAL ANALYSIS

## SUPPLEMENTAL INFORMATION

Supplemental information can be found online.

## EXPERIMENTAL MODEL AND STUDY PARTICIPANT DETAILS

### Bacterial stains and growth conditions

All *E. coli* and *P. aeruginosa* strains were grown in Lysogeny Broth (LB) medium at 37 . Where necessary, antibiotics were used at the following concentrations: for *E. coli—*chloramphenicol (25 μg ml^−1^), kanamycin (50 μg ml^−1^), gentamicin (15 μg ml^−1^), carbenicillin (100 μg ml^−1^) or tetracycline (25 μg ml^−1^); for *P. aeruginosa* — gentamicin (50 μg ml^−1^), carbenicillin (300 μg ml^−1^) or tetracycline (50 μg ml^−1^). For conjugative transfer of shuttle vectors between *E. coli*—*P. aeruginosa*, Pseudomonas Isolation Agar (PIA) supplemented with the antibiotics selective for the shuttle plasmid was used to select for transconjugants. For a list of strains used in this study, see Table S4.

## METHODDETAILS

### Plasmid construction, transformation, and strain construction

The plasmids, oligonucleotides, and synthetic genes used in this study are listed in Table S4-6. For plasmid construction, the plasmid was linearized using a restriction endonuclease (New England Biolabs), and the double-stranded DNA insert was amplified using Phanta Super-Fidelity DNA Polymerase (Vazyme). The linearized plasmid and the DNA insert were purified using the E.Z.N.A.® Gel Extraction Kit (OMEGA). Subsequently, the plasmid was assembled via Gibson assembly using the Hieff Clone® Plus Multi One Step Cloning Kit (Yeasen) and then transformed into *E. coli* DH5α for plasmid enrichment. All plasmids were verified by Sanger sequencing.

For plasmid transformation, commercial chemically competent cells were used for *E. coli* DH5α and *E. coli* BL21(DE3). Competent *E. coli* BW25113 cells were prepared using the Ultra-Competent Cell Prep Kit (Sangon Biotech). p403s and its derivative plasmids were transformed into *P. aeruginosa* using standard electroporation methods (2.5 kV, 200 Ω, 25 μF, 0.2 cm cuvette). pRSF1010, pVSG, and their derivative plasmids were transferred into *P. aeruginosa* strains via conjugation. In brief, overnight cultures of *P. aeruginosa* (recipient) and *E. coli* S17-λpir (donor) containing the plasmid were first diluted to an OD_600_ of 1. Then, the diluted recipient and donor cultures were mixed at a volume ratio of 1 mL (*P. aeruginosa*): 2 mL (*E. coli* S17-λpir). The mixed cells were centrifuged at 7,000 rpm for 2 min, and the pellet was resuspended in 100 µl of LB. The suspension was spotted onto a 0.45-μm sterile filter membrane placed on LB agar plates and incubated at 37 for 10 h. Cells from the filter membrane were collected, resuspended in 500 µl of LB, and then serially diluted. Aliquots of appropriate dilutions were plated onto *Pseudomonas* Isolation Agar (BD Difco) supplemented with corresponding antibiotics. The transformation efficiency was calculated by counting colonies and correcting for the dilution factor.

For genome editing in *E. coli* BW25113, sgRNA-donor plasmids were introduced into *E. coli* BW25113 cells harboring pECcas9 via electroporation^42^. Cells were cultured in the presence of 1% L-arabinose at 37 until the OD_600_ reached 0.4, then chilled on ice and harvested by centrifugation at 4 . Cell pellets were washed three times with ice-cold 10% glycerol and resuspended in 1 ml of ice-cold 10% glycerol. A 100 µl aliquot of the cell suspension was mixed with up to 5 µg of sgRNA-donor plasmid DNA, and the mixture was transferred to a 0.1 cm gap cuvette and electroporated using a MicroPulser Electroporator (Bio-Rad) at 1.8 kV. After electroporation, cells were incubated in 1 ml of LB medium at 37 for 1 hour, then plated on LB agar containing 50 µg/ml streptomycin sulfate. Positive clones were confirmed by PCR and sequencing, after which they were cultured in LB medium with 10% sucrose to cure the editing plasmid.

Genome editing in *P. aeruginosa* PA14 was performed using the pCasPA/pACRISPR two-plasmid system as described previously^43^. Briefly, pCasPA was electroporated into PA14. Electrocompetent cells were prepared from overnight cultures by washing with ice-cold 300 mM sucrose. After electroporation and recovery in LB at 37 for 2 h, cells were plated on LB agar with tetracycline. A single colony harboring pCasPA was then made electrocompetent and transformed with pACRISPR (carbenicillin resistant) expressing a 20-nt sgRNA and a donor template with homology arms. After electroporation, cells were recovered in LB with 10 mM L-arabinose at 30 for 2.5 h and plated on LB agar containing tetracycline, carbenicillin, and 0.5% L-arabinose. Colonies were screened by PCR and Sanger sequencing. To cure pACRISPR, verified mutants were grown at 42 without antibiotics, then plated on LB with 5% sucrose. pCasPA was subsequently cured by culturing in LB with 5% sucrose at 37 .

### Mammalian cell culture, transfection and genomic DNA extraction

HEK293T cells were cultured in Dulbecco’s Modified Eagle Medium (DMEM) supplemented with 10% fetal bovine serum (FBS) and 100 U mL ¹ penicillin-streptomycin at 37 °C with 5% CO_2_ incubation. For transfection, cells were seeded into 24-well plates (Corning) one day prior to transfection at a density of 1×10^5^ cells per well and transfected at ∼80% confluency using Hieff Clone® Liposomal 2000 Transfection Reagent (Yeasen) following the manufacture’s recommended protocol. For each well, cells were co-transfected with 350 ng of ErCas12a plasmid and 150 ng of crRNA or CracrVA1 plasmid. Twenty-four hours after transfection, the medium was replaced with fresh medium containing puromycin at a final concentration of 4 µg/mL, and the cells were incubated for additional 3 days. For deep sequencing analysis, cells were washed with PBS, and genomic DNA was extracted by adding 40 µL of lysis buffer (Thermo Fisher Scientific) supplemented with 25 µg mL ¹ proteinase K (Thermo Fisher Scientific). The mixture was incubated at 55 °C for 5 min, followed by inactivation at 98 °C for 5 min.

### High throughput sequencing

The genomic region flanking the CRISPR target site for each gene in HEK293T cells was amplified using the Platinum Direct PCR Universal Master Mix (Thermo Fisher Scientific). PCR primers were designed to generate amplicons of 180-250 bp covering the regions of interest. Unique barcodes were incorporated into the forward and reverse ends of the PCR products to enable sample multiplexing for high throughput sequencing. Following amplification, equal amounts of barcoded products were pooled and purified using the E.Z.N.A. Gel Extraction Kit (Omega). Purified libraries were subjected to commercial sequencing on the NovaSeq platform (GENEWIZ). Raw sequencing reads were demultiplexed based on barcode sequences and analyzed according to previously described method^44^. Editing efficiency was calculated as the proportion of sequencing reads containing insertions or deletions (indels) relative to total reads.

### Phage propagation

Infections with bacteriophage T7 were carried out in LB medium supplemented with 10 mM MgCl . A single plaque was picked and purified twice to ensure clonal homogeneity. Subsequently, 5 ml of bacterial culture at an OD of 0.1 was infected with phage at a multiplicity of infection (MOI) of 0.1, and the infection was allowed to proceed overnight. The resulting lysate was centrifuged at 4,000 g for 20 min at 4°C, and the supernatant was filtered through a 0.22-μm pore-size syringe filter.

### Phage infection assays

To prepare plates for phage spotting assays, 150 μl of the bacterial culture (OD_600_ reached 0.8-1.0) was mixed with 3 mL of molten 0.5% LB agar (supplemented with 10 mM MgSO4 and corresponding antibiotics), and the mixture was poured over an 1.2% LB agar plates. After the top agar solidified, 2.0 μl drops of 10-fold serial dilutions of the phage reservoir were spotted on the surface. The plates were incubated at 37 for 8-12 h. Plaques were counted and the plaque forming units per milliliter (PFU/ml) were calculated. For each tested phage lysate, the phage titer was quantified as PFU/ml. The PFU count was determined based on the lowest phage dilution spot that allowed individual quantification of PFUs. Using the dilution factor of that spot, the PFU value per milliliter was calculated. When a vague clearing zone, rather than distinguishable individual plaques, was observed, the lowest phage concentration at which this phenomenon occurred was recorded as 10 plaques. If no clearing zone was observed, the highest dilution spot was considered to represent 1 plaque, which was taken as the detection limit of the assay. Each experiment included 3-4 independent biological replicates.

### Dilution plating assay

For bacterial plating assays, individual colonies were inoculated into LB broth containing appropriate antibiotic(s), and bacterial cultures were allowed to grow overnight to stationary phase. 10-fold serial dilutions of these cultures were prepared with LB broth, and 2.0 μl of these dilutions were spotted on LB agar plates (supplemented with appropriate antibiotics, 0.3% *L*-arabinose or 0.5% glucose as needed). Colonies were counted after overnight incubation to calculate colony formation units per milliliter (CFU/ml). Each experiment included three independent biological replicates.

### RNA isolation

Overnight cultures were inoculated (at a ratio of 1:100) into 10 mL of LB medium supplemented with appropriate antibiotics and then grown at 37°C until they reached the logarithmic phase. 2 mL of bacterial culture was subjected to centrifugation (7000 rpm for 2 min at 4) to collect the cells, then the total RNA was extracted using the E.Z.N.A. Bacterial RNA Kit (Omega), following the standard guidelines. DNA contaminants were removed using DNase (New England Biolabs) according to the manufacture’s instruction. RNA concentration was determined using a NanoDrop One Spectrophotometer (Thermo Fisher Scientific).

### 5**′** RACE analysis

5′ RACE was performed using the template-switching enzyme (New England Biolabs). In brief, 1 μg of the total RNA was mixed with 1 mM specific reverse primer (listed in Table S6), and the mixture was incubated at 85 for 5 min and then immediately placed on ice. Subsequently, a template-switching reverse-transcription reaction was performed to generate complementary DNA (cDNA) with a universal sequence of choice on their 3′ end (introduced by a template switching oligonucleotide). The cDNA products were subjected to PCR reactions using another specific primer and a primer against the template-switching oligonucleotide (TSO). The PCR products were analyzed on a gel and verified by Sanger sequencing. If double peaks were observed, the PCR products were cloned into a vector. Following transformation, 10 randomly picked clones were PCR-amplified and re-sequenced by Sanger sequencing.

### Small RNA sequencing and analysis

A total of 50 μg of RNA was treated with T4 polynucleotide kinase (New England Biolabs) according to the manufacture’s protocol. The RNA was purified using the phenol-chloroform method and then precipitated with an equal volume of isopropanol and 0.1 volume of 3M sodium acetate. The RNA molecules ranging from 30 to 300 nt were selected to construct a small RNA library with the NEXTFLEX Small RNA-Seq Kit (Bioo Scientific) and then subjected to Illumina HiSeq sequencing (paired-end, 150-bp reads). The raw data was processed to remove adapters. The resulting reads were mapped to the *cracr*, *creR* or *creA* sequence using custom Perl scripts^1^.

### RT-qPCR

cDNA was synthesized using the M-MuLV Reverse Transcriptase (NEB) and quantified using the ChamQ Blue Universal SYBR qPCR Master Mix (Vazyme) on an Applied Biosystems ViiA^TM^ 7 Real-Time PCR System. Three biological samples were analyzed for each experimental setting, and for each sample, three technical replicates were included. The primers are listed in Table S6.

### Fluorescence measurement

To measure fluorescence in *P. aeruginosa*, cells containing *mCherry* were cultured to the late logarithmic growth phase. Then, 200 µL of each culture was transferred to a microplate for the simultaneous measurement of OD_600_ and mCherry fluorescence using the Synergy H4 Hybrid multimode microplate reader (BioTek). For each of the three biological replicates, the ratio of fluorescence to OD_600_ was determined, and the mean and standard deviation were calculated.

### Northern blot

RNA sample was mixed with an equal volume of RNA Gel Loading Dye (Thermo Fisher Scientific), heated to 85 for 5 min and then cooled on ice for 2 min. Then, the RNA samples were resolved by denaturing 10% TBE-Urea PAGE and then electrotransferred onto Biodyne B nylon membranes (Pall). Membranes were UV-crosslinked, then pre-hybridized 1 hour at 42, followed by the addition of biotin-labelled ssDNA probe (listed in Table S6) and incubation overnight at 42 . The signal was detected using the Chemiluminescent Nucleic Acid Detection Module Kit (Thermo Fisher Scientific), according to the manufacturer’s protocol. The membrane was imaged using the Tanon 5200 Multi chemiluminescent imaging system (Tanon Science & Technology). Each blotting assay was repeated using separate biological samples, with a representative result provided.

### Western blot

Overnight cultures were inoculated 1:100 into 250 mL of LB medium supplemented with appropriate antibiotics and grown at 37 °C to logarithmic phase (OD_600_ ∼0.8). Cells were harvested by centrifugation at 12,000×g for 2 min. The pellet was resuspended in lysis buffer (20 mM HEPES pH 7.5, 500 mM NaCl, 5% glycerol, 1 mM DTT) and sonicated. Cell debris was removed by centrifugation at 12,000×g for 1 h. Total protein (5 mg) was separated by 12% SDS PAGE and transferred onto a nitrocellulose membrane (Millipore). A parallel gel was stained with Coomassie Blue as a loading control. The membrane was blocked with 5% non fat milk in TBST (20 mM Tris HCl pH 7.5, 150 mM NaCl, 0.1% Tween 20) for 1 h at room temperature, then incubated with 6x-His Tag Monoclonal Antibody (1:2000, Thermo Fisher Scientific) overnight at 4 °C. After three washes with TBST, the membrane was incubated with Goat anti-Mouse IgG (H+L) secondary antibody, HRP (1:5000, Thermo Fisher Scientific) for 1 h at room temperature. Protein bands were visualized using enhanced chemiluminescence (ECL) substrate (Bio Rad) and imaged with a chemiluminescence imaging system (Tanon 5200).

### Protein expression and purification

Plasmids expressing Csy1-Csy4 and crRNA/CracrIF2/CreA2 were co-transformed into *E. coli* BL21(DE3) cells. Protein expression was induced by 0.2 mM IPTG when the cell density reached an OD_600_ of 0.8. Cells were cultured at 18°C for 12 hours, harvested by centrifugation, and resuspended in lysis buffer (50 mM Tris-HCl pH 8.0, 300 mM NaCl, 10 mM imidazole, 1 mM PMSF). Protein extraction was performed via sonication, followed by centrifugation at 20,000 × g for 50 minutes at 4°C to remove cellular debris.

The clarified supernatant was subjected to affinity purification using a 5 mL HisTrap™ FF (Cytiva) column equilibrated in wash buffer containing 50 mM Tris-HCl pH 8.0, 300 mM NaCl, 30 mM imidazole, and eluted using a gradient against elution buffer (lysis buffer containing 500 mM imidazole). Elution fractions were concentrated and further purified using a Superdex-200 Increase 10/300 GL (GE Healthcare) column equilibrated with a buffer containing 10 mM Tris-HCl pH 8.0, 200 mM NaCl, 5 mM dithiothreitol. The purified protein was analyzed by SDS-PAGE.

For Acr F24 purification, the open reading frame was cloned into pGEX-6P-1 plasmid for fusion expression with GST-tag, and subsequently transferred into *E. coli* BL21(DE3) cells. GST-Acr F24 expression was induced with 0.1 mM IPTG upon reaching an OD of 0.8, followed by cultivation at 18°C for 12 hours. The purification procedure is the same as Csy complex except that the GSTrap™ HP (Cytiva) column, lysis buffer (20 mM Hepes pH 7.5, 500 mM NaCl, 1 mM PMSF) and elution buffer (20 mM Hepes pH 7.5, 500 mM NaCl, 10 mM reduced glutathione, 1mM dithiothreitol). The fusion protein was then digested with PreScission protease at 4 overnight to remove GST-tag.

For Cas12a purification, the *cas12a* gene was cloned into the pET28a plasmid to fuse a 6×His-SUMO tag, and the resulting plasmid was subsequently transferred into *E. coli* BL21(DE3) cells. Growth medium plus inoculum was grown at 37 until the cell density reached an OD_600_ of 0.4; the temperature was then decreased to 22 . Growth was continued until OD_600_ reached 0.8, when 0.2 mM IPTG (final concentration) was added to induce 6×His-SUMO-Cas12a expression. The culture was induced for 18 h before cells were harvested and frozen at -80 until purification. The cell pellet was resuspended in lysis buffer (20 mM HEPES pH 7.5, 500 mM NaCl, 5 mM MgCl, 20 mM imidazole) supplemented with protease inhibitors (Roche cOmplete, EDTA-free). Protein extraction was performed via sonication, followed by centrifugation at 20,000 × g for 1 h at 4 to remove cellular debris. The lysate was filtered through 0.22 μm filters (Millipore Stericup) and applied to a 5 ml HisTrap™ FF nickel column (Cytiva), washed, and then eluted with a gradient of imidazole. Fractions containing protein of the expected size were pooled, concentrated, and further purified using a Superdex-200 Increase 10/300 GL (GE Healthcare) column equilibrated with buffer containing 10 mM Tris-HCl pH 8.0, 200 mM NaCl, 5 mM dithiothreitol. The purified protein was analyzed by SDS-PAGE. The fusion protein was then digested with SUMO protease (Beyotime, P2312S) at 4 overnight to remove the 6×His-SUMO tag.

### Small RNA synthesis

For synthesis of crRNA, CracrVA1, and the corresponding mutant RNAs, ss DNA oligonucleotides (Table S6) and their complementary strands were synthesized by Genewiz. The two complementary ssDNA were mixed, heated to 98, and then allowed to cool naturally to room temperature to anneal, generating double-stranded transcription templates. *In vitro* transcription was performed using the HyperScribe™ T7 High Yield RNA Synthesis Kit (APExBIO, K1047-25) according to the manufacturer’s instructions to produce crRNA, CracrVA1, and the mutant RNAs. After transcription, the template DNA was removed by DNase I treatment, and the RNA was purified by phenol-chloroform extraction.

### DNA cleavage assays

Cas12a-mediated cleavage assays were performed in cleavage buffer containing 20 mM HEPES (pH 7.5), 150 mM KCl, 10 mM MgCl, 1% glycerol and 0.5 mM DTT. 2 μM Cas12a was pre-assembled with either 3 μM CracrVA1, CracrVA1mut or crRNA at 37 for 10 min. To generate P*_cas12a_* dsDNA substrates, the FAM-labelled forward strand was heated to 98 in the presence of a twofold excess of unlabeled complementary strand and allowed to cool slowly to room temperature. The cleavage reaction was initiated by adding 20 nM P*_cas12a_* dsDNA to the pre assembled Cas12a complexes (at final concentrations of 500, 1000, and 2000 nM), and the mixture was incubated at 37 for 30 min. Reactions were quenched with formamide-containing dye and analyzed on 15% urea-PAGE. The gels were visualized using a Tanon-5200 Multi instrument (Tanon Science & Technology Co. Ltd., Shanghai, China).

### Electrophoretic mobility shift assay (EMSA)

The oligonucleotides used for preparing dsDNA substrates are listed in Table S6, and dsDNA substrate preparation was performed as described above. For Figure 1E, Cas12a (2 μM) was assembled with crRNA, CracrVA1, or its mutant RNAs (3 μM) in binding buffer (20 mM HEPES, 150 mM NaCl, 5% glycerol and 0.5 mM DTT, pH 7.5) at 37 for 10 min. For binding assays, reaction mixtures containing 20 nM FAM-labeled P*_cas12a_* dsDNA and varying concentrations (0, 125, 250, 500, or 1000 nM) of pre-assembled Cas12a-CracrVA1, Cas12a-CracrVA1mut, or Cas12a-crRNA complexes were incubated at 37 for 15 min. For Figure 4E, DNA binding reactions were performed in binding buffer at 37 for 15 min. Each reaction contained non target DNA (0, 0.15, 0.3125, 0.625, or 1.25 μM), FAM labeled target P*_creT2_* DNA (0.04 μM), and 0.8 μM of either Csy crRNA, Csy CreA2, or Csy CracrIF2 complex, with or without AcrIF24 protein (9.6 μM). Following incubation, samples were resolved on a 6% native polyacrylamide gel. Free dsDNA and shifted complexes were detected using the Tanon 5200 Multi System (Tanon Science & Technology Co., Ltd., Shanghai, China).

### GST pull-down assay

The GST-Acr F24 fusion protein was incubated with Csy-crRNA, Csy-CreA2 and Csy-Cracr2, respectively, according to the molecular weight of 1:2 with Mag-MK GST Fusion Protein Purification beads (BBI: C650031). At the same time, only cascade with magnetic beads were used as a negative control. The mixture was gently incubated at 4 for 3h and then the supernatant was discarded. Pre-cooled wash buffer (20 mM Hepes pH 7.5, 150 mM NaCl, 1% Triton X-100) was added incrementally along the wall of the tube to resuspend the beads, followed by four sequential wash cycles. Finally, elution buffer (20 mM Hepes pH 7.5, 500 mM NaCl, 10 mM reduced glutathione, 1 mM dithiothreitol) was applied to elute bound complexes. Proteins were analyzed by SDS-PAGE to verify interactions.

### Cryo-EM sample preparation and data acquisition

4 μL aliquots of Csy proteins complexed with crRNA/Cracr2 /CreA2 at a concentration of 4/4.5/1.5 mg/mL were respectively applied to discharged 300-mesh Quantifoil R1.2/1.3 grids (Quantifoil, Micro Tools GmbH, Germany). Grids were blotted for 4.5 s and plunged into liquid ethane using an FEI Mark IV Vitrobot operated at 8 and 100% humidity.

Cryo-EM data were collected on a 300 kV Titan Krios G3 equipped with a Gatan K3 detector and a GIF Quantum energy filter (slit width of 20 eV). The defocus values ranged from -1.3 µm to -1.8 µm. Each stack of 32 frames was exposed for 2.56 s, and the exposure time of each frame was 0.08 s. The micrographs were automatically collected with the AutoEMation program^45^ in super-resolution counting mode with a binned pixel size of 0.8374 Å. The total dose of each stack was about 50 e^-^ Å^-2^. The alignment and summation of all 32 frames in each stack were performed using the whole-image motion correction program MotionCor2^46^.

### Cryo-EM data processing

All cryo-EM data were processed using cryoSPARC^47^. Contrast transfer function (CTF) parameters were estimated using patch-CTF. Blob picker was used for initial templates generation via 2D classification. Template picker was then used for all particle-picking tasks. For the cascade-crRNA, 732,730 particles were extracted from 928 micrographs. Subsequent two-dimensional (2D) classification, multi-class ab-initio reconstruction, nonuniform refinement and local refinement of the best class were performed, resulting in the resolution of 3.11 Å. Following the same workflow, 2,226,978 particles were extracted from 2,954 micrographs and 1,246,776 particles were extracted from 2,388 micrographs. Subsequently, 146,897 particles were used to resolve the structure of cascade-Cracr2 at 2.90 Å resolution, and 122,663 particles were used to resolve the structure of cascade-CreA at 3.32 Å resolution, respectively. All structures were determined with C1 symmetry. Resolution is reported using the gold-standard Fourier shell correlation with 0.143 cutoff. See Figure S11-13 for protein Cryo-EM data processing flow. See Table S8 for data collection, refinement and validation statistics.

### Cryo-EM model building and refinement

The initial models were derived using AlphaFold3^48^. We assigned predicted models into our density maps by UCSF ChimeraX^49^. The models were further optimized by Coot^50^. The final models of all datasets were refined against the corresponding maps using PHENIX^51^ in real space with secondary structure and geometry restraints. The structures were validated through examination of the Clash scores, Molprobity scores and statistics of the Ramachandran plots by PHENIX (Table S8). All the figures were created in PyMOL and UCSF ChimeraX.

**Search for *cracr* genes**

The *creA* or *creR* sequences were used as queries for BLASTN (v2.16.0+) under the blastn-short mode against the NCBI RefSeq, IMG/VR and IMG/PR databases. The searches were performed with the following parameters: -word_size 7 -evalue 10 -outfmt 6. Raw BLAST outputs were stringently filtered: only hits with an alignment length ≥ 20 bp and covering the characteristic *cracr* region (query start ≤ 25, query end ≥ 26) were retained; short and non-specific matches were discarded. Genomic coordinates of candidate target sequences were calculated from the BLAST results, strand orientation was corrected, and redundant sites were removed. Full-length phage *cracr* homolog sequences were extracted using Seqkit v2 to construct a candidate dataset. To eliminate interference from native CRISPR-Cas repeat sequences, BLAST hits adjacent to Cas genes were filtered based on genome annotation files, and only valid targets located in non-Cas regions were kept. Genomic coordinates were recalibrated, and putative *cracr* sequences were subsequently extracted.

A hidden Markov model (HMM) profile of AcrIF24 was used to search for homologous genes in the NCBI RefSeq, IMG/VR and IMG/PR databases. For each identified homolog, intergenic regions longer than 40 bp located within 1.5 kb upstream and downstream were extracted. These intergenic regions were then subjected to BLASTn searches using known Cracr/CreR sequences as queries with the following parameters: -reward 1 -penalty -2 -gapopen 5 -gapextend 2 -word_size 7 -dust no -evalue 10. More CracrIF candidates were subsequently identified by manual inspection of repeat like sequences.

### Hierarchical clustering of *cracr* repeat regions

The repeat regions derived from *cracr* sequences with clear target loci were used for hierarchical clustering analysis. Pairwise p-distances were calculated based on global alignment using the following parameters: match score = 2, mismatch score = –1, gap open penalty = –0.5, and gap extension penalty = –0.1. For each sequence pair, the optimal alignment was used, and the p-distance was computed as p = 1 – m/L, where m is the number of matched sites and L is the total alignment length. A symmetric distance matrix was then constructed. Hierarchical clustering was performed using the complete linkage method based on this distance matrix. The resulting dendrogram was converted into Newick format, with sequence IDs and branch lengths annotated at leaf nodes and topological relationships at internal nodes. The Newick tree file was visualized and annotated using iTOL (https://itol.embl.de/upload.cgi).

### Phylogenetic analysis of AcrIF24 proteins

The AcrIF24 HMM profile was used to search the NCBI RefSeq, IMG/VR and IMG/PR databases. Hits with at least 90% sequence coverage were retained. Amino acid sequences were aligned using MAFFT (v7.525) with the FFT-NS-1000 algorithm. A phylogenetic tree was constructed from the alignment using FastTree (v2.1.11) under the WAG evolutionary model with gamma-distributed site rates. The tree was visualized and annotated using iTOL (https://itol.embl.de/upload.cgi).

## QUANTIFICATION AND STATISTICAL ANALYSIS

For all statistical analyses, experiments were performed with at least three independent replicates (as indicated in the figure legends). *P* values were calculated using two-sided Student’s *t* test unless otherwise stated. All other results were derived from at least two independent experiments with consistent observations

## Supplemental information

**Figure S1.**
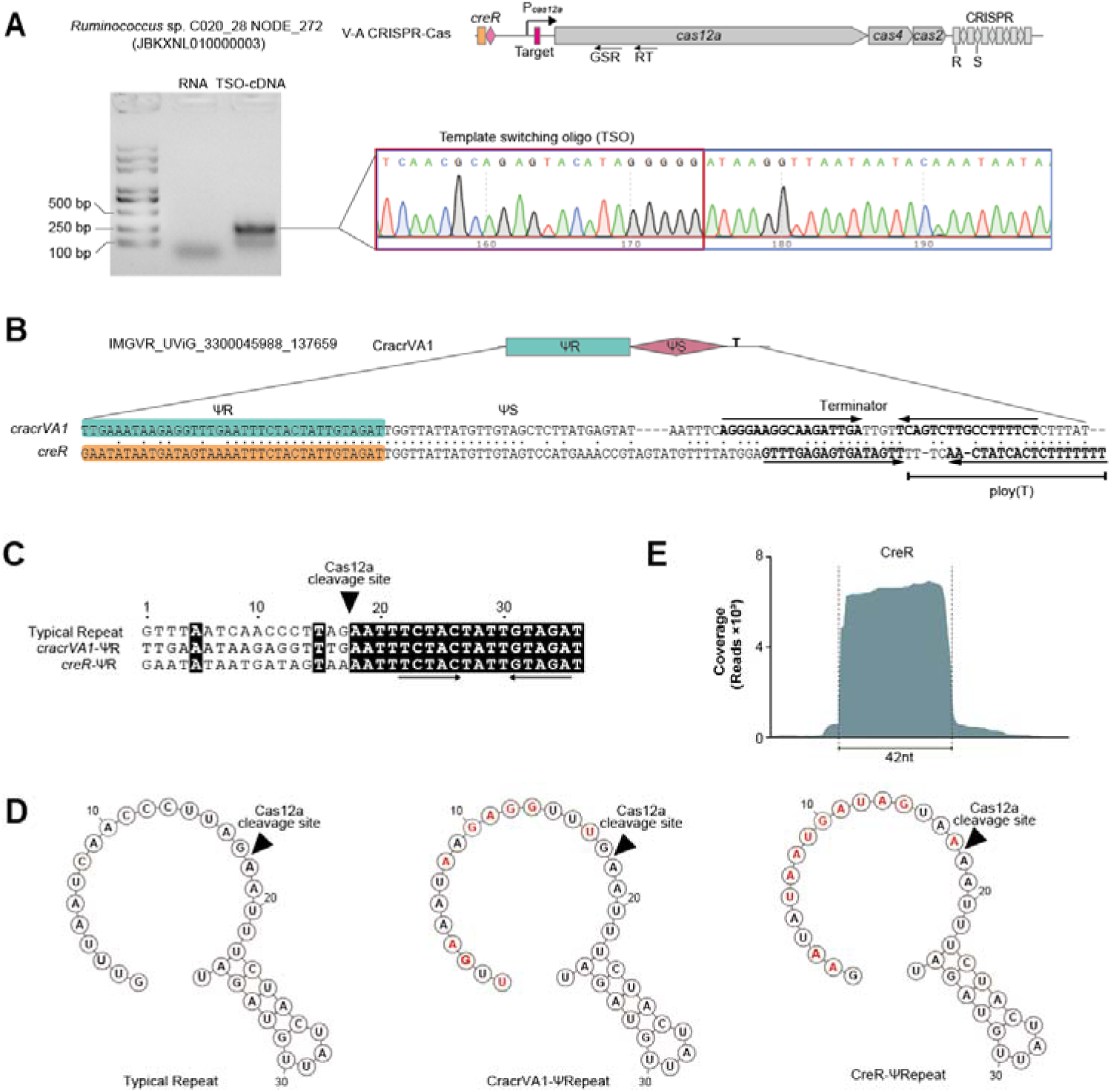
CracrVA1 and its target CRISPR-Cas system, related to. Figure 1. (A) 5′ RACE analysis of the *Ruminococcus* V-A CRISPR-Cas locus, which contains a CreR RNA gene mediating autorepression. The *cas12a*-specific primers used for reverse transcription (RT) and RACE PCR (GSR) are indicated. DNase-treated RNA samples served as the negative control. PCR products were subjected to Sanger sequencing to determine their 5′ ends. cDNA, complementary DNA. TSO, template switching oligo. (B) Full sequence of the investigated *cracrVA1* gene and its sequence similarity to the cognate *creR* gene (dots indicate identical nucleotides). The rho-independent terminator comprises an RNA hairpin (indicated by inverted arrows) followed by consecutive thymines. Palindromic nucleotides are shown in bold. (C) Alignment of the ΨR sequences to typical CRISPR repeat shown in panel A. Palindromic nucleotides are indicated with inverted arrows, and the Cas12a cleavage site on RNA products is marked. (D) Predicted RNA structure of the ΨR sequences. Nucleotides that differ from typical CRISPR repeat are highlighted in red. (E) Small RNA-seq data of CreR. Small RNA was extracted from *E. coli* cells expressing codon-optimized *Ruminococcus* sp. C020_28 Cas12a (under the control of its native promoter) along with the *creR* gene.

**Figure S2.**
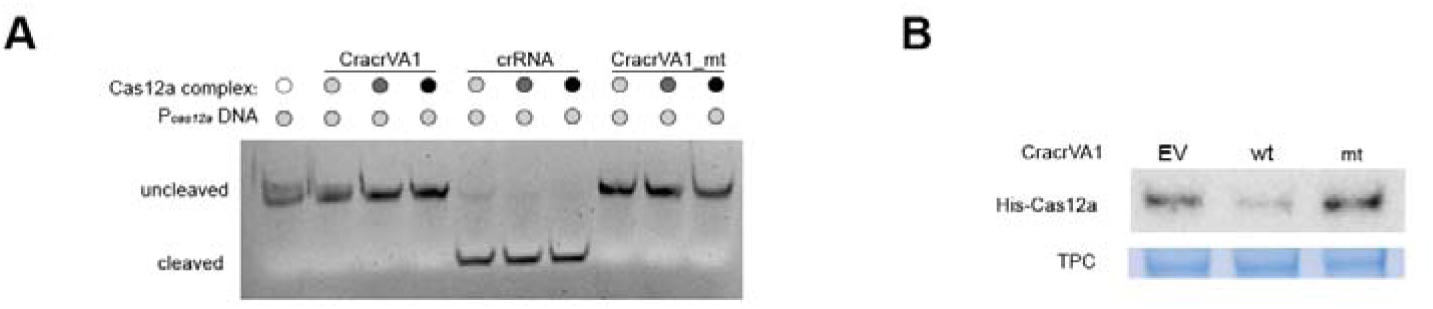
Nuclease assay and Western blotting of Cas12a, related to Figure 1. (A) Cleavage of P*_cas12_* DNA by Cas12a complexes guided by a targeting crRNA, CracVA1, or its spacer-mutant (mt) at different concentrations. (B) Western blot analysis of Cas12a in *E. coli* BW25113 cells harboring the type V-A locus from *Ruminococcus* sp. C020_28 (on the chromosome) following introduction of an empty vector (EV) or its derivatives expressing wild-type (wt) or spacer-mutated (mt) CracrVA1. Coomassie blue staining (bottom) serves as a loading control for total protein (TPC).

**Figure S3.**
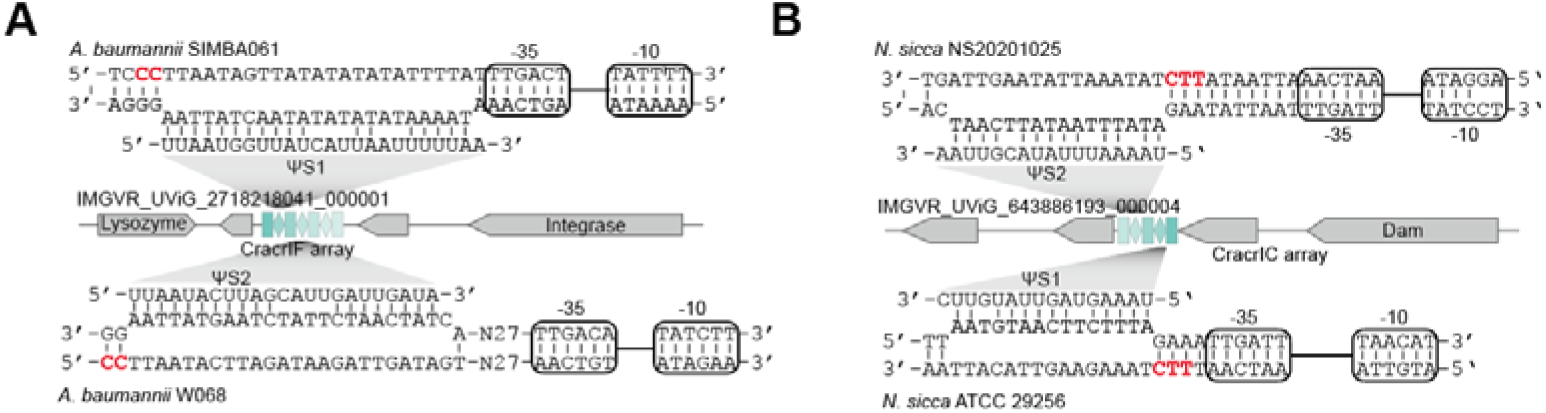
Representative Cracr arrays encoding two or more viral crlRNA species, and their base pairings to predicted target CRISPR-Cas loci from different bacterial strains. Predicted promoter elements (−35 and −10) located within the CRISPR-Cas loci are indicated. PAM nucleotides are highlighted in red, related to Figure 2.

**Figure S4.**
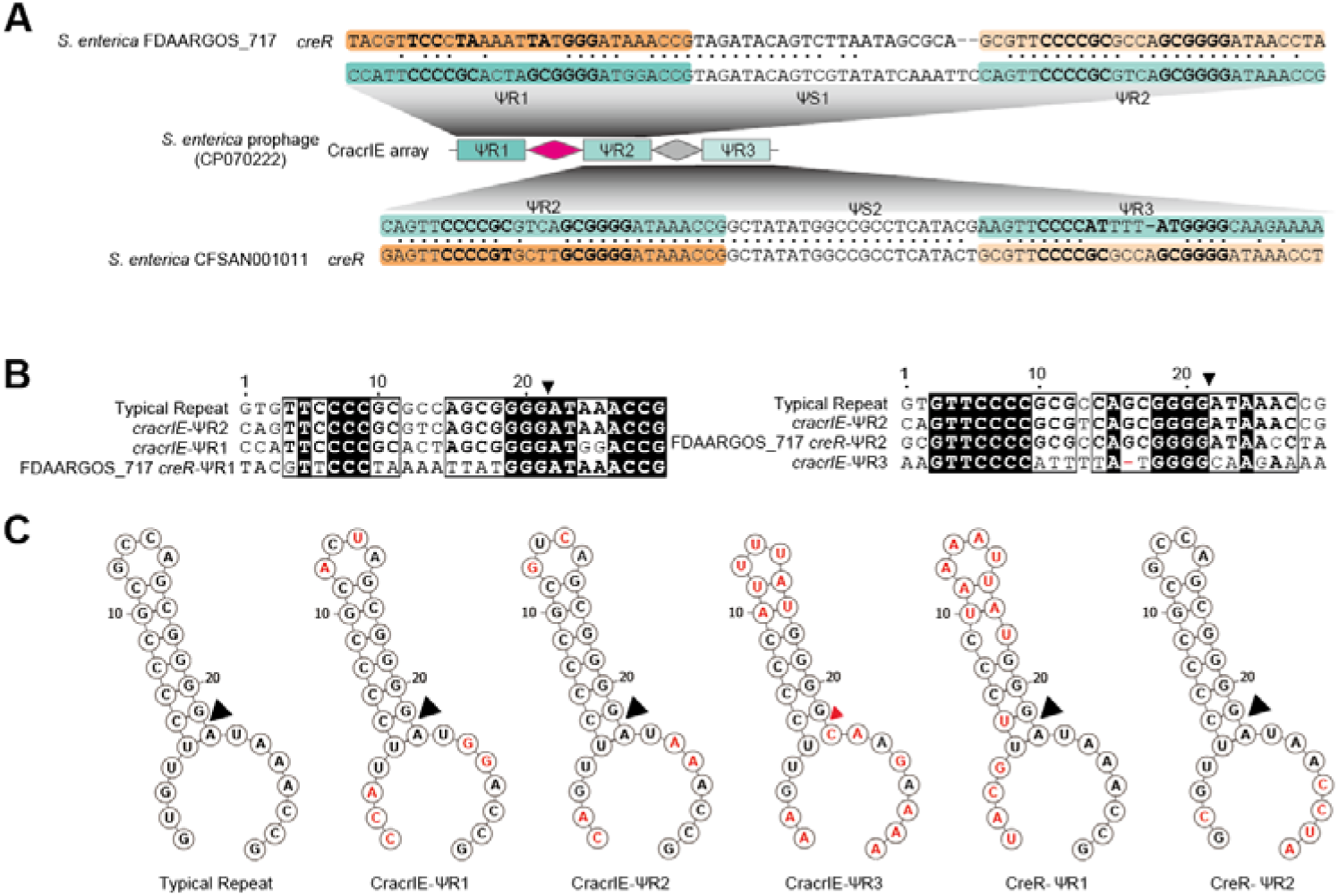
Bioinformatic analysis of an CracrIE array carried by a *S. enterica* prophage, related to Figure 2. (A) Sequence of the investigated CracrIE array and its sequence similarity to each *creR* element associated with the two targeted *S. enterica* CRISPR-Cas loci (dots indicate identical nucleotides). Palindromic nucleotides are shown in bold. (B) Alignment of ΨR sequences located 5′ upstream (left) or 3′ downstream (right) of each ‘spacer’ sequence to the typical CRISPR repeat. A red hyphen in the ΨR3 sequence denotes a single-nucleotide deletion. (C) Predicted RNA structures of each ΨR sequence. Nucleotides that differ from the typical CRISPR repeat are highlighted in red. Black and red (less efficient) triangles indicate the Cas6 cleavage sites on mature RNA products.

**Figure S5.**
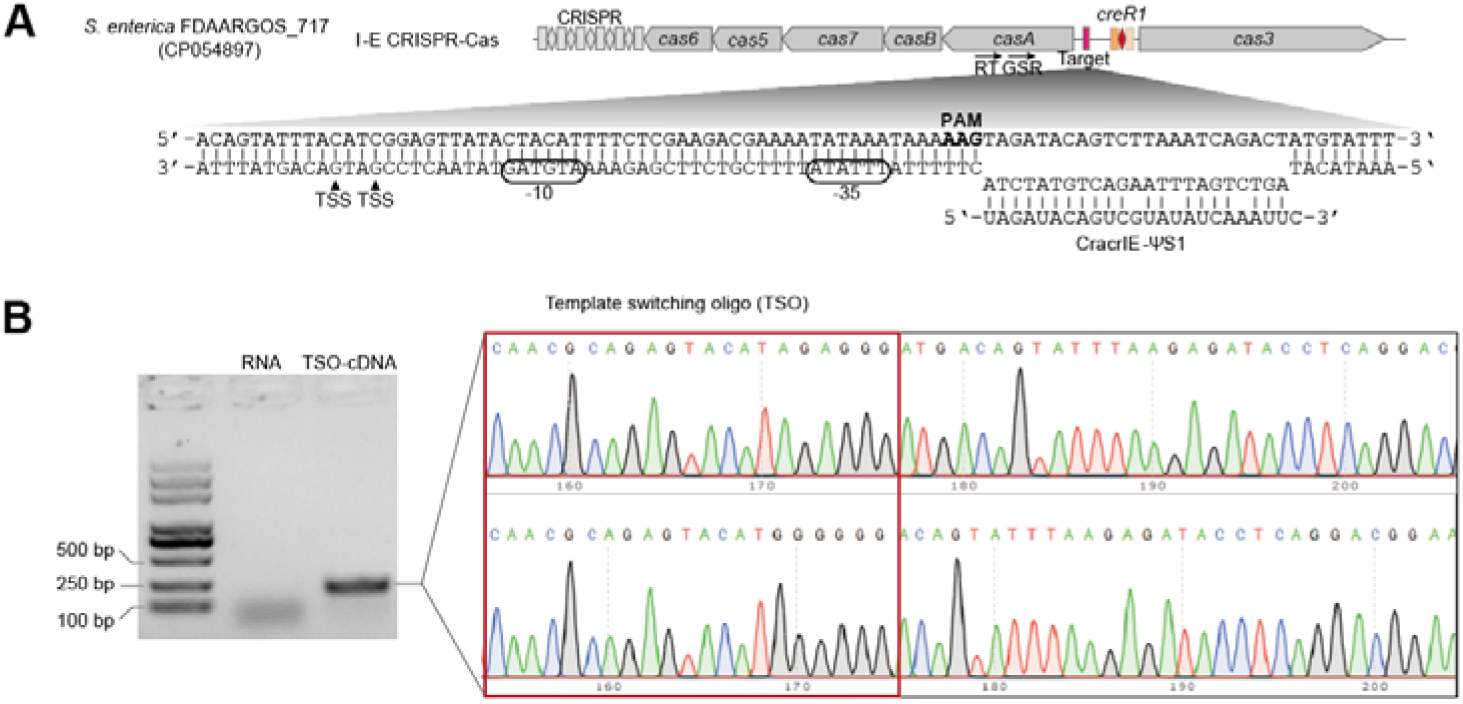
Characterization of the *casA* promoter of *S. enterica* FDAARGOS_717, related to Figure 2. (A) Schematic of the *S. enterica* CRISPR-Cas locus targeted by the first ΨS of the investigated CracrIE array. TSS, transcription start site. Predicted promoter elements −10 and −35 are indicated. (B) 5′ RACE analysis. The *casA*-specific primers used for reverse transcription (RT) and RACE PCR (GSR) are indicated in panel A. DNase-treated RNA samples served as the negative control. PCR products were subjected to Sanger sequencing to determine their 5′ ends. cDNA, complementary DNA.

**Figure S6.**
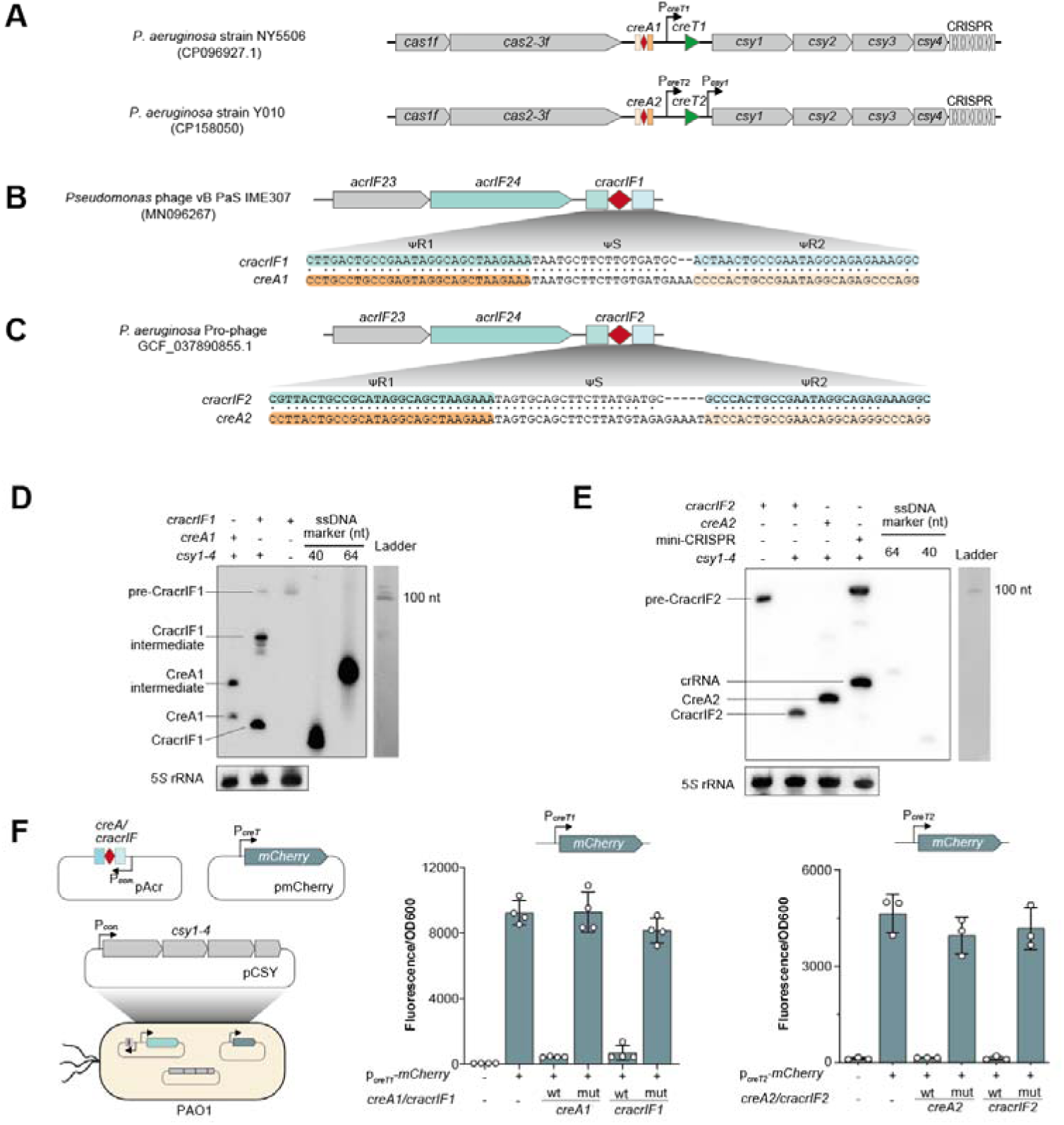
Characterization of CracrIF RNAs in *P. aeruginosa*, related to Figure 3. (A) Schematic of two example *P. aeruginosa* CRISPR-Cas loci containing the CreTA1 or CreTA2 module. (B, C), Schematics of two representative I-F CracrIFs (CracrIF1 and CracrIF2) that show similarity to CreA1 and CreA2, respectively. Dots indicate identical nucleotides. (D, E) Northern blot analyses of mature CracrIF and CreA RNAs. These RNA molecules were expressed in PAO1 cells expressing the PA14 Csy proteins (Csy1-4). PAO1 cells lacking Csy proteins (producing precursor RNAs) were included as controls. 5*S* rRNA served as the internal control. Biotin-labeled single-stranded oligos (40 nt or 64 nt) served as size markers. A single-stranded RNA ladder was co-electrophoresized. (F) Assessment of the transcription-inhibiting activity of CracrIF and CreA RNAs using an mCherry reporter assay. Both wild-type (wt) and mutated (mut; the first three nucleotides of ΨS were subjected to mutation) versions of the these regulatory RNAs were tested. Error bars, mean±s.d. (n=3).

**Figure S7.**
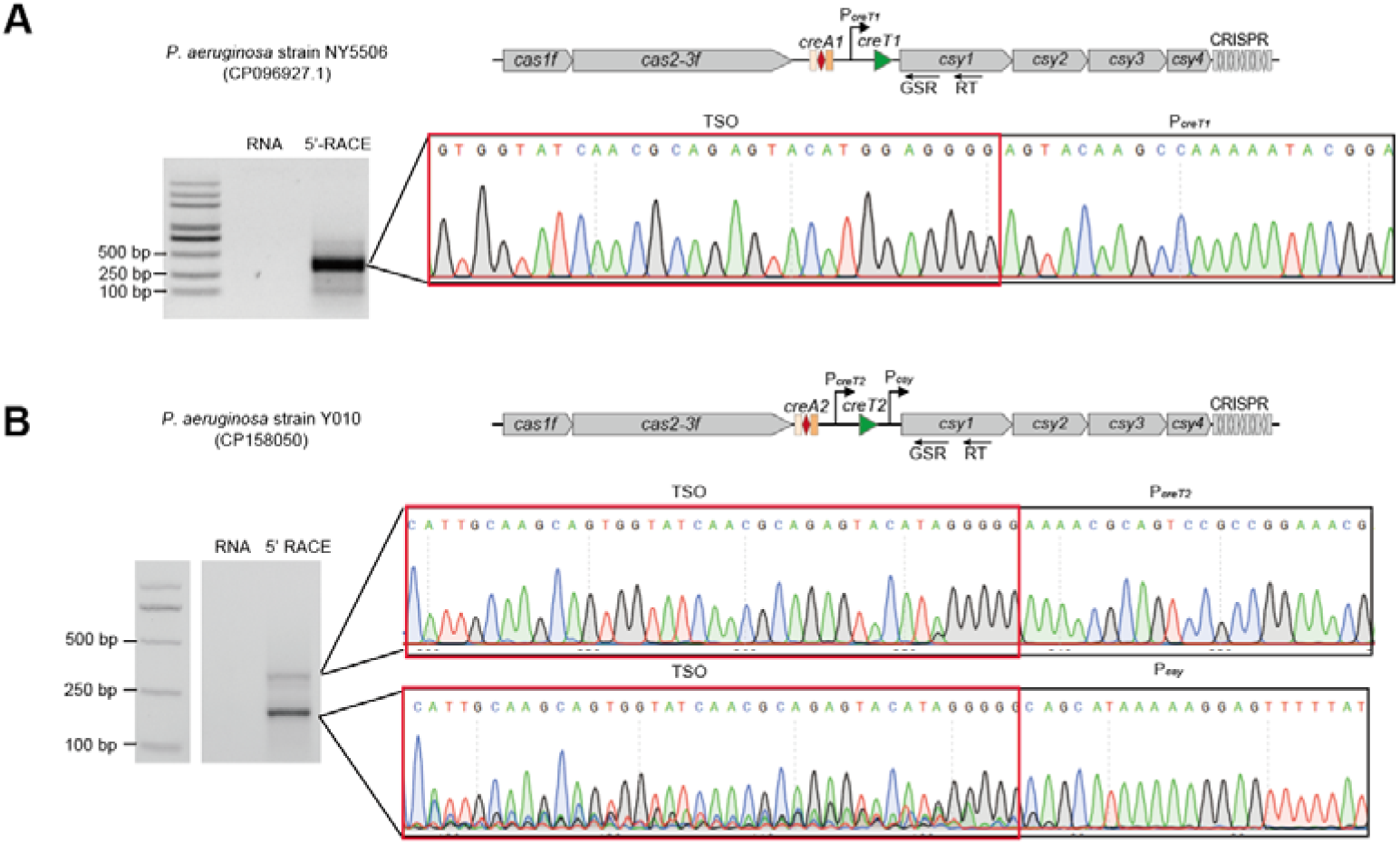
5′ RACE analysis of the *csy* transcripts from *P. aeruginosa* NY5506 (a) and Y010 (b) CRISPR-Cas loci, related to Figure 3. The *csy1*-specific primers used for reverse transcription (RT) and RACE PCR (GSR) are indicated. DNase-treated RNA samples served as the negative control. PCR products were subjected to Sanger sequencing to determine their 5′ ends. cDNA, complementary DNA.

**Figure S8.**
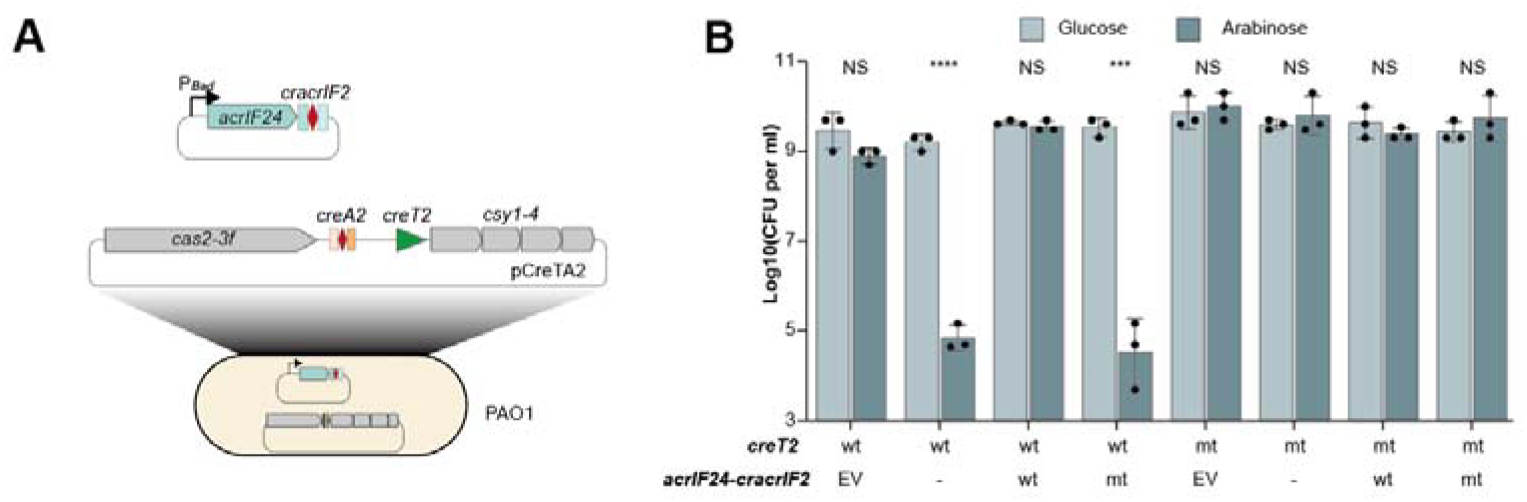
Experimental design and results of the PAO1 dilution spotting assay, related to Figure 3. (A) Schematic of the experimental design. PAO1 cells were engineered to harbor a plasmid-born Y010 *cas* operon (containing the CreTA2 module) and transformed with another plasmid carrying a P*_BAD_*-driven *acrIF24-cracrIF2* cassette. (B) Dilution spotting of PAO1 cells on arabinose (inducing) or glucose (repressing) medium. EV, empty vector. The *acrIF24* was introduced alone (-) or together with either wild-type (wt) or spacer-mutated (mt) *cracrIF2*. An inactive *creT2* mutant serves as a negative control. CFU, colony forming units. Error bars, mean±s.d. (n=3). *P* values from two-sided Student’s *t* test. ***, *P*≤0.001; ****, *P*≤0.0001; NS, not significant.

**Figure S9.**
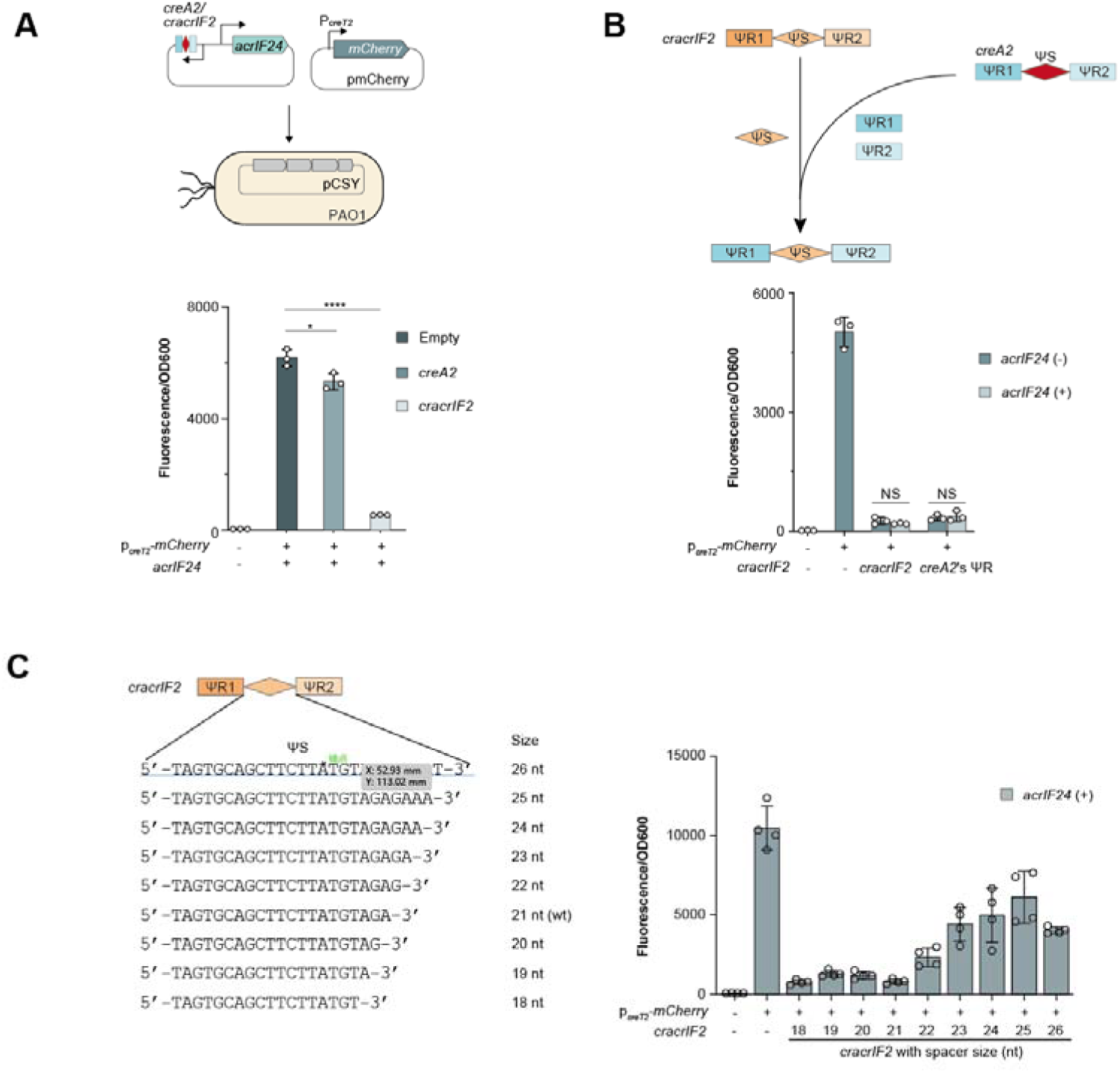
Assessing the effect of AcrIF24 on the regulatory function of CracrIF2- and CreA2-guided Csy complex, related to Figure 4. (A) The experimental design and the data showing the inhibitory effects of CracrIF2 or CreA2 on P*_creT2_* in the presence of AcrIF24. (B) The inhibitory effects of CracrIF2 before and after replacing its ΨR sequences with the counterparts from CreA2. Error bars, mean±s.d. (n=3). *P* values from two-sided Student’s *t* test. *, *P*≤0.05; ****, *P*≤0.0001; NS, not significant. (C) Impact of spacer size on the regulatory activity of CracrIF2 RNAs in the presence of AcrIF24. Error bars, mean±s.d. (n=4).

**Figure S10.**
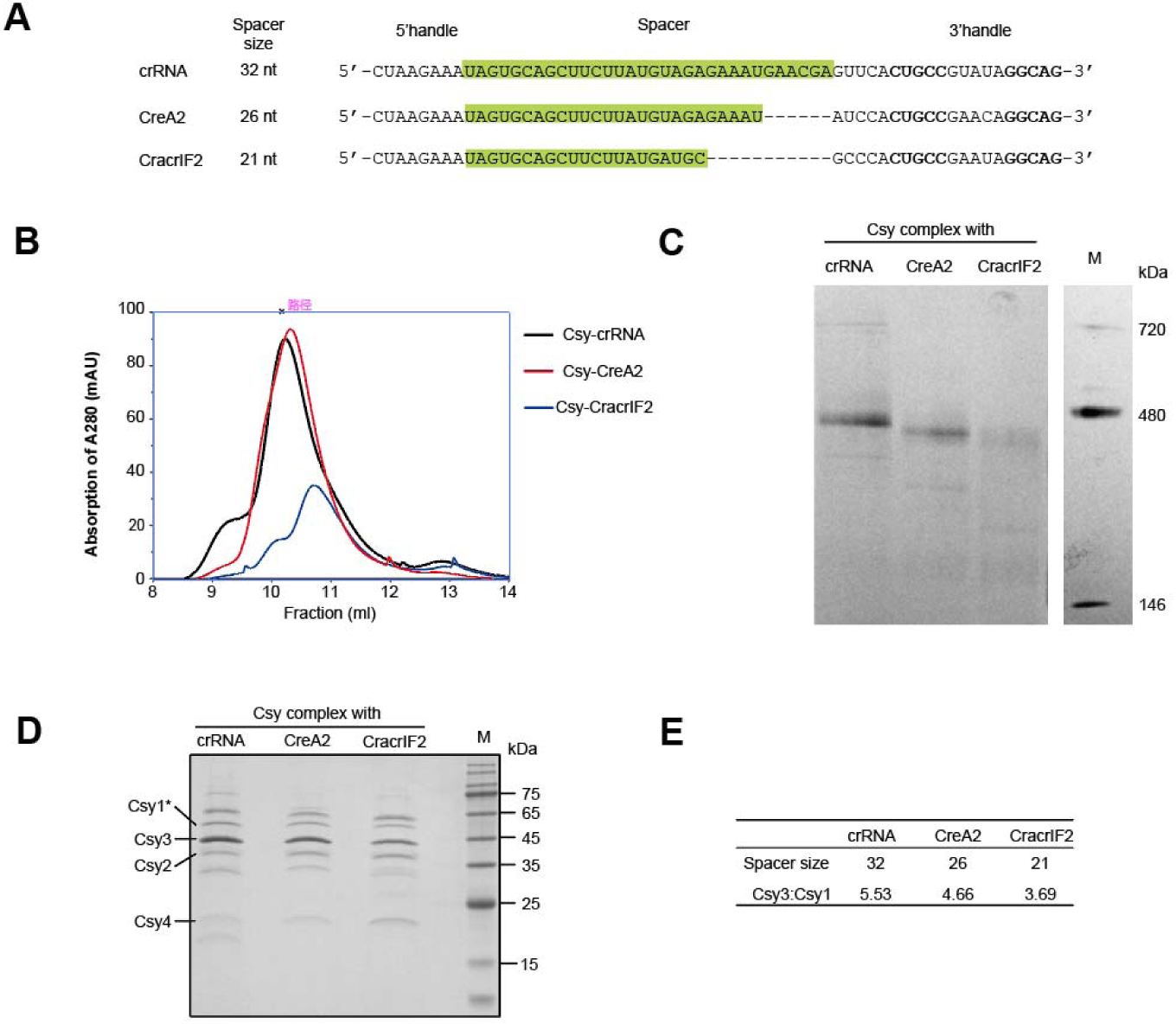
Comparative analysis of CracrIF2, CreA2, and a canonical crRNA in complex with Csy proteins, related to Figure 4. (A) Nucleotide sequences of CreA2, CracrIF2, and a canonical crRNA (designed based on CreA2 with five additional spacer nucleotides that are not complementary to P*_creT2_*, resulting a typical spacer size of 32 nt). (B) Size exclusion chromatography (SEC) traces showing co-expressed Csy1-4 proteins (with His*6 tagged Csy1) with crRNA, CreA2, or CracrIF2 in *E. coli*. mAU, milliabsorbance units. (C) Native PAGE analysis of Csy complexes bound to CracrIF2, CreA2, or the cognate crRNA. Csy proteins and guide RNAs were co-expressed in *E. coli* BL21(DE3) and subsequently co-purified for analysis. (D) SDS-PAGE analysis of peak fractions purified by SEC (see panel B). (E) Molar ratio of Csy3 and Csy1 estimated based on band intensities shown in panel D and their molecular weights.

**Figure S11.**
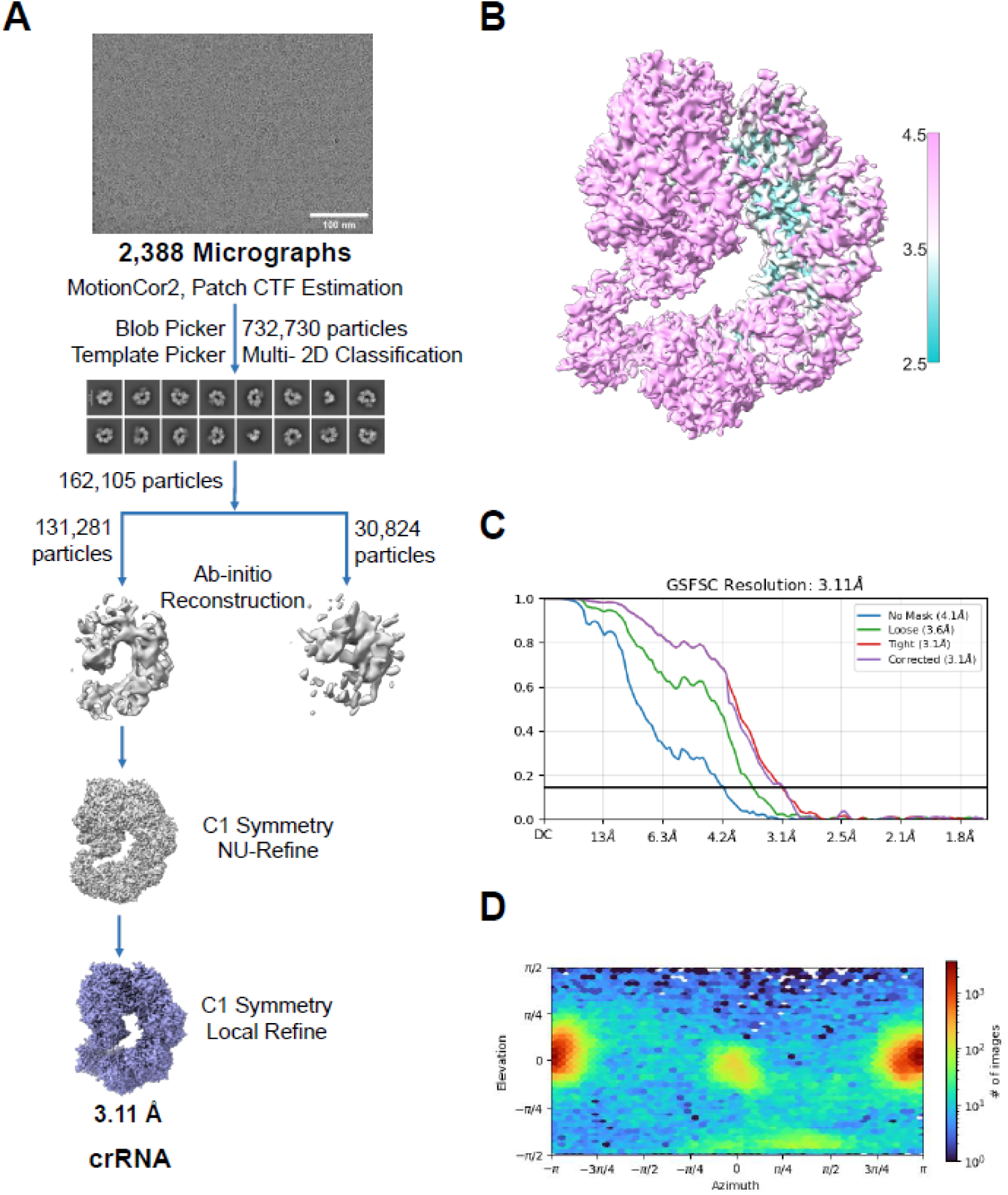
Data processing of Csy-crRNA dataset, related to Figure 4. (A) The representative micrograph, 2D class averages and the flowchart for data processing of crRNA. (B-D) The local resolution, gold-standard Fourier shell correlation (FSC) curves from 3D refinement, and the angular distribution of particles used in the final reconstructions of crRNA.

**Figure S12.**
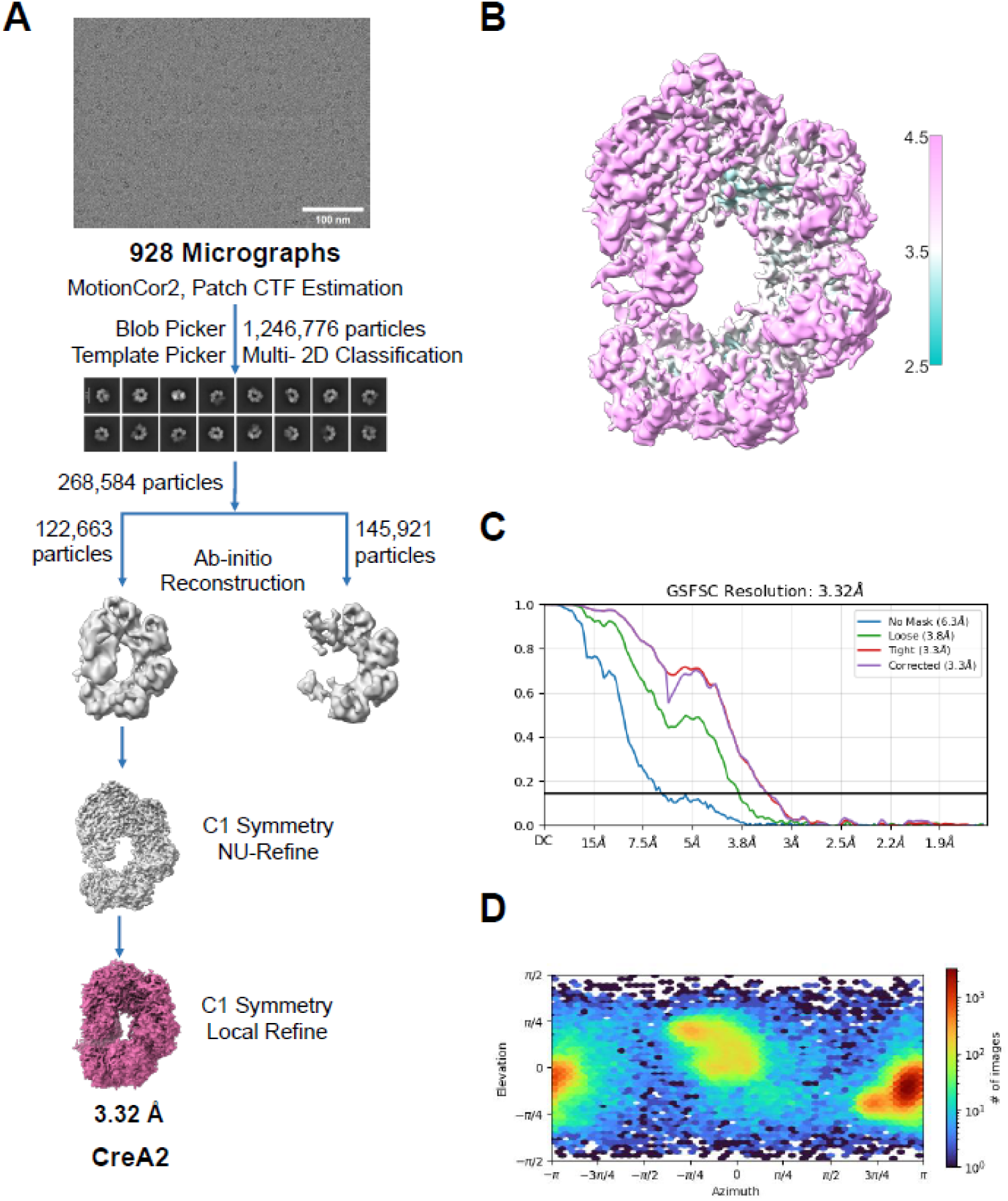
Data processing of Csy-CreA2 dataset, related to Figure 4. (A) The representative micrograph, 2D class averages and the flowchart for data processing of CreA2. (B-D) The local resolution, gold-standard Fourier shell correlation (FSC) curves from 3D refinement, and the angular distribution of particles used in the final reconstructions of CreA2.

**Figure S13.**
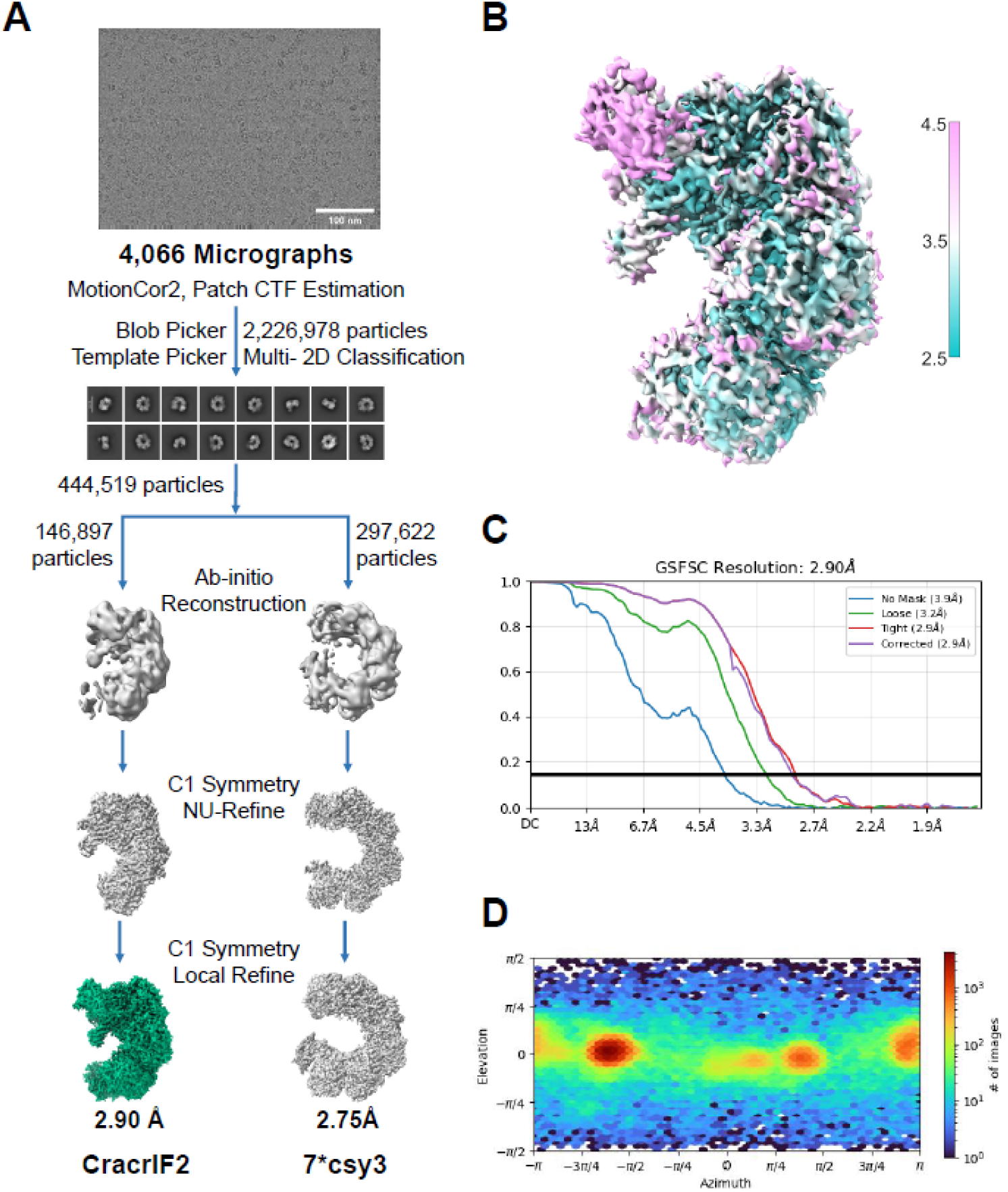
Data processing of Csy-CracrIF2 dataset, related to Figure 4. (A) The representative micrograph, 2D class averages and the flowchart for data processing of CracrIF2. (B-D) The local resolution, gold-standard Fourier shell correlation (FSC) curves from 3D refinement, and the angular distribution of particles used in the final reconstructions of CracrIF2.

**Figure S14.**
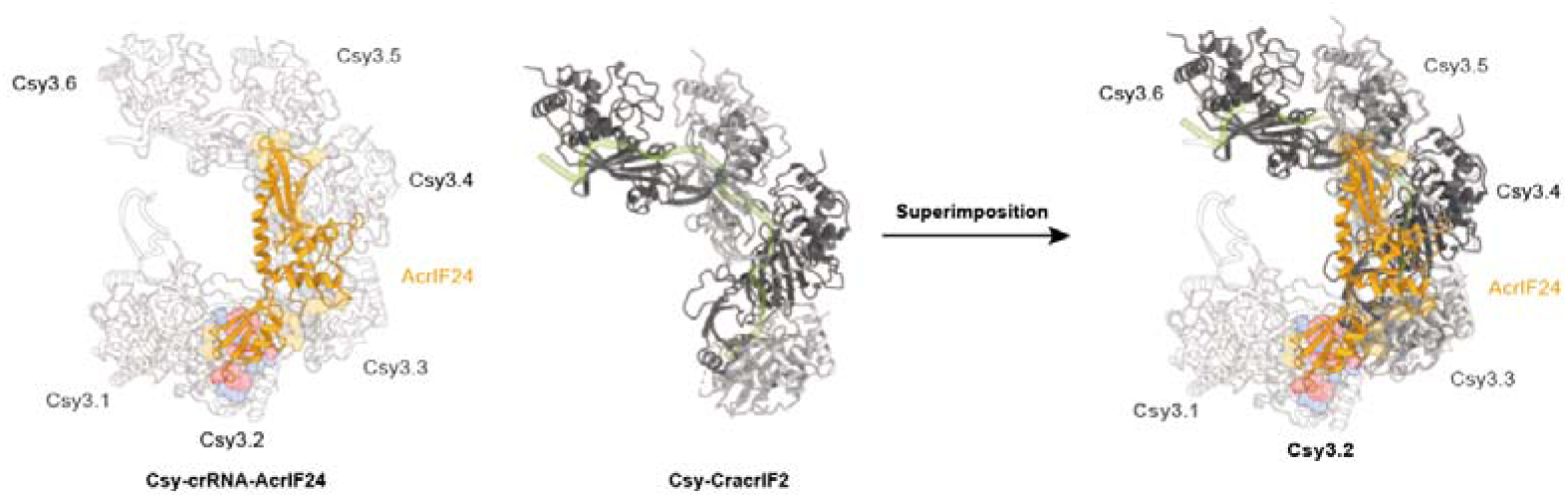
Superimposition of the Csy-CracrIF2 structure onto the previously reported Csy-crRNA-AcrIF24 structure (PDB: 7ELN)^21^, related to Figure 4. Note that only the Csy3 backbone subunits were displayed. The interacting residues between AcrIF24 (colored yellow) and the Csy3 subunits (colored grey) are depicted in the surface model. The residues of AcrIF24 and Csy3.2 that are involved in their interface are highlighted in red and blue, respectively.

**Figure S15.**
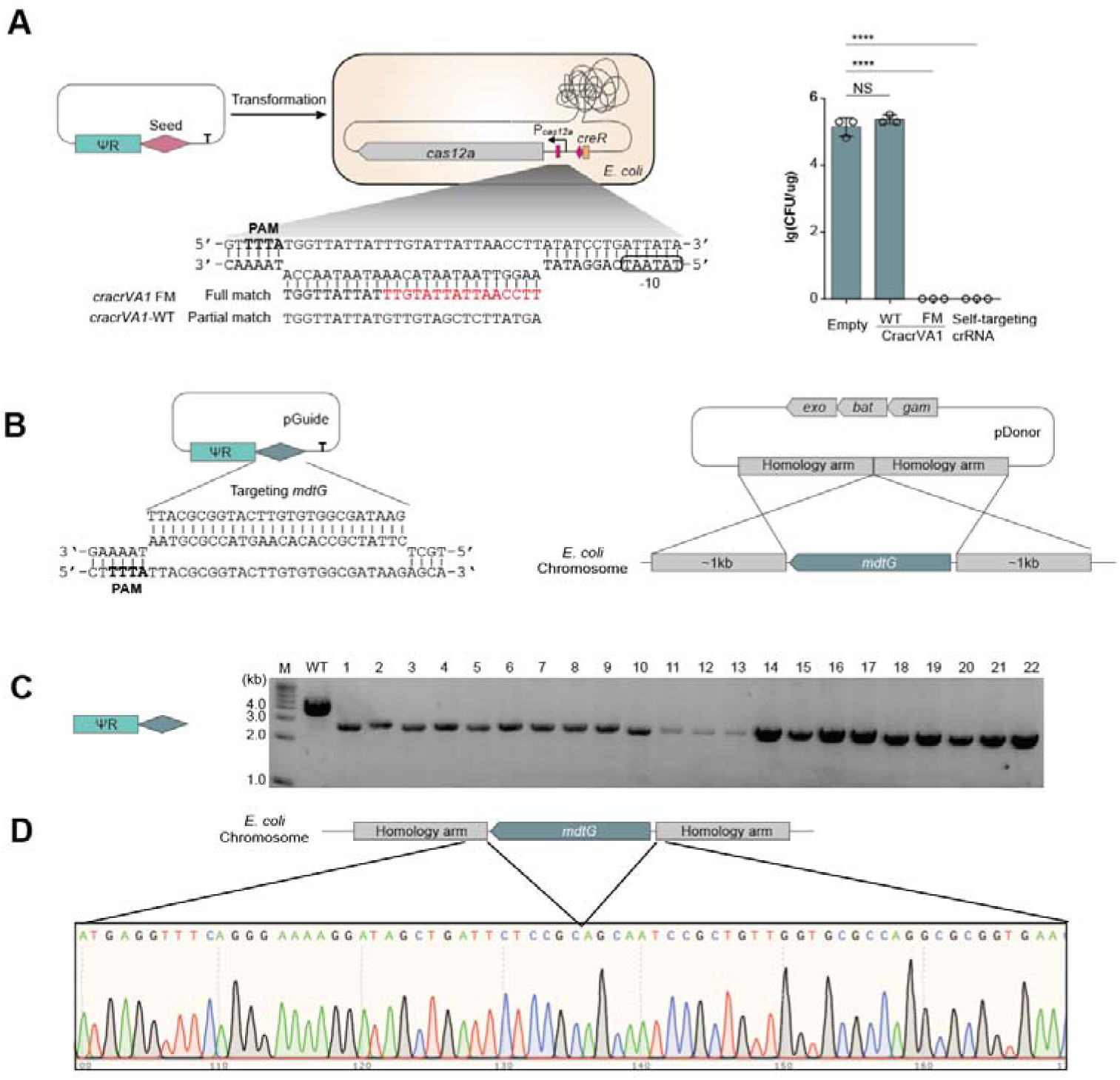
Repurposing CracrVA1 to edit the *E. coli* genome, related to Figure 1. (A) Transformation efficiency of *E. coli* cells harboring *Ruminococcus* sp. C020_28 *cas12a* and its native promoter, using a plasmid expressing variant Cracrs that fully match P*_cas12a_*. A typical CRISPR array encoding a self-targeting crRNA served as a control. Error bars, mean±s.d. (n=3). *P* values from two-sided Student’s *t* test. *, *P*≤0.05; NS, not significant. (B) Schematic of the editing plasmids (pGuide and pDonor) designed to knock out the non-essential *mdtG* gene from the *E. coli* chromosome. pDonor carries the λ-Red recombination system (*exo*, *bat*, and *gam*) along with two homology arms derived from genomic sequences flanking *mdtG*. (C) PCR analysis showing the genotypes of 20 randomly selected edited clones. (D) Sanger sequencing result of a representative edited clone.

**Figure S16.**
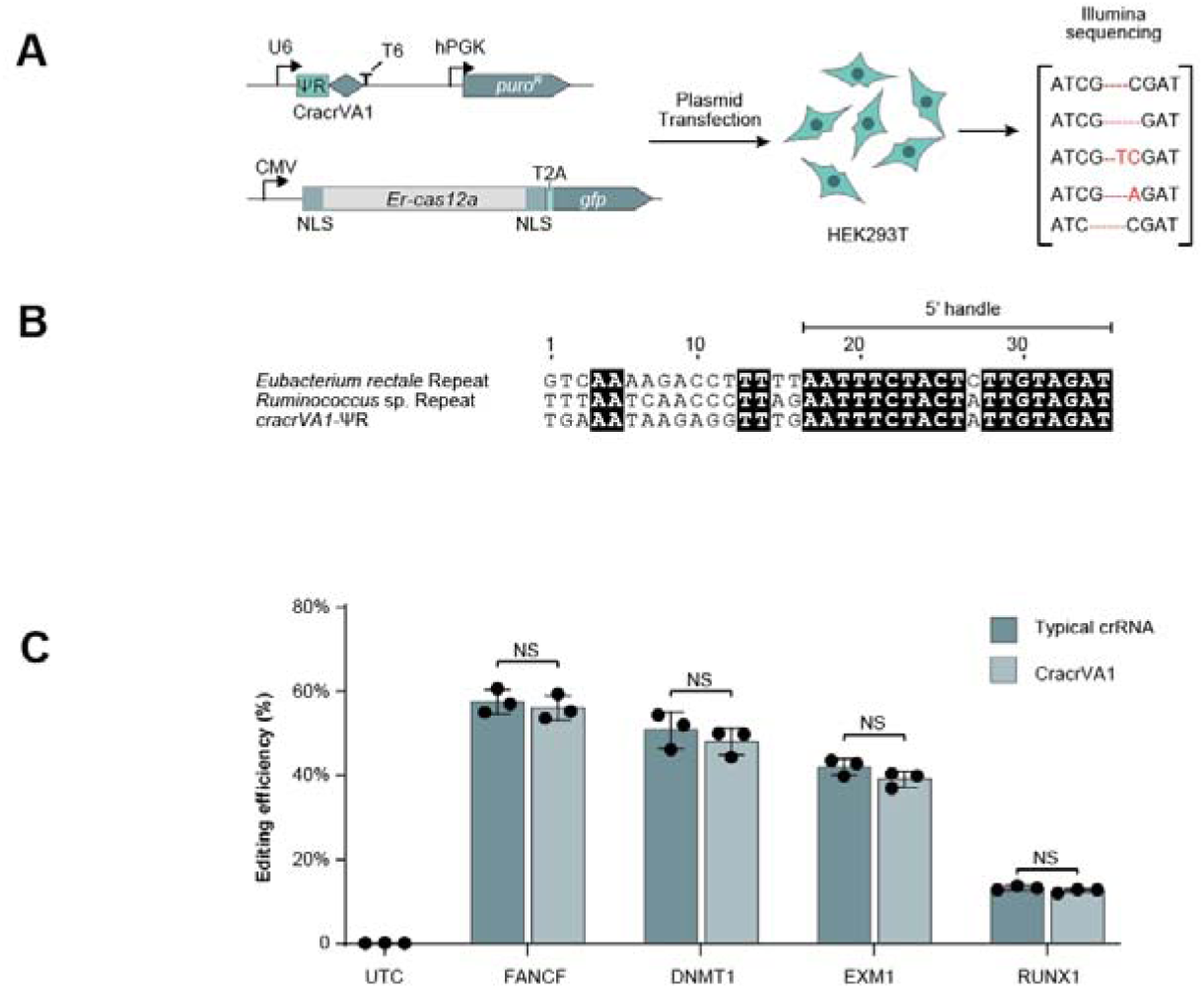
Engineering of the CracrVA1 RNA for Cas12a-based mammalian genome editing, related to Figure 1. (A) The experimental design. ErCas12a, *Eubacterium rectale* Cas12a. NLS, nuclear localization signal. The eukaryotic U6 promoter and T6 terminator were used to control guide RNA expression. UTC, untransfected control. (B) Sequence similarity between Cracr ΨR and the typical CRISPR repeat sequences of *Ruminococcus* sp. C020_28 and *Eubacterium rectale* Cas12a. Nucleotides encoding the 5′ handle on mature RNAs are indicated. (C) Editing efficiency (indel frequency) of four human genes by ErCas12a complexed with either typical crRNA guides or reprogrammed CracrVA1 RNAs. UTC, untransfected control. Error bars, mean±s.d. (n=3). *P* values from two-sided Student’s *t* test. **, *P*≤0.01; ***, *P*≤0.001; ****, *P*≤0.0001; NS, not significant.

## Notes

### Competing Interest Statement

The authors have declared no competing interest.

### Summary of Updates

This version of the manuscript has been revised to update the following. The abstract has been rephrased for improved clarity and conciseness. The Results section has been substantially expanded with three key additions: functional validation of Cracr activity in type V‑A and I‑E CRISPR‑Cas systems, experimental evidence for Cracr as an effective single‑guide RNA in gene editing, and a comprehensive bioinformatic analysis to identify Cracr candidates across diverse microbial genomes. The author list has been reordered according to contributions, and three new co‑authors have been included: Junyuan Xue, Lei Cai, and Eugene V. Koonin. Despite these extensive changes, the main theme and core conclusions of the study remain unchanged.

